# Naturally occurring combinations of receptors from single cell transcriptomics in endothelial cells

**DOI:** 10.1101/2021.01.04.425317

**Authors:** Sergii Domanskyi, Alex Hakansson, Michelle Meng, Benjamin K Pham, Joshua S. Graff Zivin, Carlo Piermarocchi, Giovanni Paternostro, Napoleone Ferrara

## Abstract

VEGF inhibitor drugs have been successful, especially in ophthalmology, but not all patients respond to them. Combinations of drugs are likely to be needed for more effective therapies of angiogenesis-related diseases. In this paper we describe naturally occurring combinations of receptors in endothelial cells that might help to understand how cells communicate and to identify targets for drug combinations. We also develop and share a new software tool called DECNEO to identify them.

Single-cell gene expression data are used to identify a set of co-expressed endothelial cell receptors, conserved among species (mice and humans) and enriched, within a network, of connections to up-regulated genes. This set includes several receptors previously shown to play a role in angiogenesis. Multiple statistical tests from large datasets, including an independent validation set, support the reproducibility, evolutionary conservation and role in angiogenesis of these naturally occurring combinations of receptors. We also show tissue-specific combinations and, in the case of choroid endothelial cells, consistency with both well-established and recent experimental findings, presented in a separate paper.

The results and methods presented here advance the understanding of signaling to endothelial cells. The methods are generally applicable to the decoding of intercellular combinations of signals.

## INTRODUCTION

Angiogenesis is the development of new blood vessels. It is a physiological process required for embryonic development, adult vascular homeostasis and tissue repair [1]. It is a fundamental process of human biology involved in many diseases, including cancer, eye diseases, arthritis, psoriasis and many other ischemic, inflammatory, infectious, and immune disorders.

During tumor progression, the new vessels provide neoplastic tissues with nutrients and oxygen; in intraocular vascular disorders, the growth of abnormal, leaky blood vessels may destroy the retina and lead to blindness [1,2]. Extensive efforts to dissect the molecular basis of angiogenesis and to identify therapeutic targets resulted in the discovery of key signaling pathways involved in vascular development and differentiation [1,3]. In particular, numerous studies have established the pivotal role of the VEGF (Vascular Endothelial Growth Factor) pathway in physiological and pathological angiogenesis and therapies targeting this pathway have achieved success in the treatment of cancer and ocular disorders [4,5]. VEGF inhibitors are currently the main class of FDA-approved anti-angiogenic drugs and they are used as a therapy for cancer and for eye diseases leading to blindness. One of us has provided fundamental contributions to the understanding of the role of VEGF in angiogenesis and to the development of these drugs [5].

VEGF inhibitor drugs have been successful, especially in ophthalmology, but not all patients respond to them [1,4,5].

Additionally, while anti-VEGF therapies have had a major impact, the use of VEGF or other angiogenic factors, including FGF1, FGF2, FGF4, VEGF-C and HGF, in order to promote collateral vessel growth and endothelial survival in patients with coronary or limb ischemia has not been clinically successful [6,5,4,7].

These observations support the hypothesis that VEGF is only part of the signal for normal vessels growth and endothelial survival. Multiple extracellular factors have been shown to influence angiogenesis in biological models and combinations of drugs acting on these molecules are likely to be needed for more effective therapies of angiogenesis-related diseases [1,4,5].

Several methods are currently used to discover novel drug combinations [8]. Beyond empirical combinations of individually effective drugs, they include the use of drug screens, using brute force (that is, for small sets, exhaustive testing of all possible combinations) or search algorithms depending on the phenotypic response [9]. They also include modeling of biological systems based on statistical associations or direct modeling of molecular interactions [10]. These approaches can also be combined, but our knowledge of the details of *in vivo* biological systems is still incomplete and progress to the clinic is therefore slow and inefficient. Understanding the regulation and role of co-expressed gene sets has been a key focus of biological investigations, starting from the discovery of the *lac* operon in bacteria by Jacobs and Monod [11]. In the case of receptors, which are both receivers of natural signals and drug targets, understanding their physiological role can be a useful guide to combination therapy development.

In this paper we describe naturally occurring combinations of genes or their products relevant for intercellular communication to endothelial cells, An increasing number of single-cell RNA-seq datasets provide information on organ and tissue-specific signaling. Dedicated initiatives such as the Human Cell Atlas, the NIH Human Biomolecular Atlas Program and the EU Lifetime Initiative [12,13] aim to map all the cell types of the human body and can inform the understanding of co-expression at the single-cell level. Co-expression at the single-cell level gives a more detailed understanding of biological regulation compared to bulk measurements.

The evolutionary age and rate of change of genes can provide information about their functions and interactions [14]. We also expect that any functionally important co-expression of genes would be conserved during evolution. Considerations analogous to these have guided combinatorial interventions before, for example the finding by Shinya Yamanaka and his collaborators that delivering the same four transcription factors can reprogram cells to a pluripotent state in both mice and humans [15].

Furthermore, while incomplete, network models of intracellular interactions are being developed using an increasing number of datasets and can contribute to the validation of new findings [10]. Endothelial cells, the main targets of VEGF, are the key cell type in angiogenesis and their receptors integrate multiple signals to decide if angiogenesis should be activated [16,6]. We show how to identify receptors linked, within network models, to significantly up-regulated genes by calculating a measure called network enrichment. It is a measure of statistical association, but, in some cases, it might represent downstream changes due to the signals transmitted by the receptors.

We show here that an integrated analysis of multiple single cell RNA-seq datasets, using co-expression, level of expression, evolutionary conservation and network enrichment can be used to identify naturally occurring combinations of receptors in endothelial cells. This set includes VEGF receptors and other receptors previously shown to play a role in angiogenesis. Figures 1 and 2 provide a visual introduction to our analysis. We then show that these naturally occurring combinations are in part tissue-specific and that, in the case of choroid endothelial cells, results are consistent with both well-established and recent experimental findings, presented in a separate paper [17].

**Figure 1.**
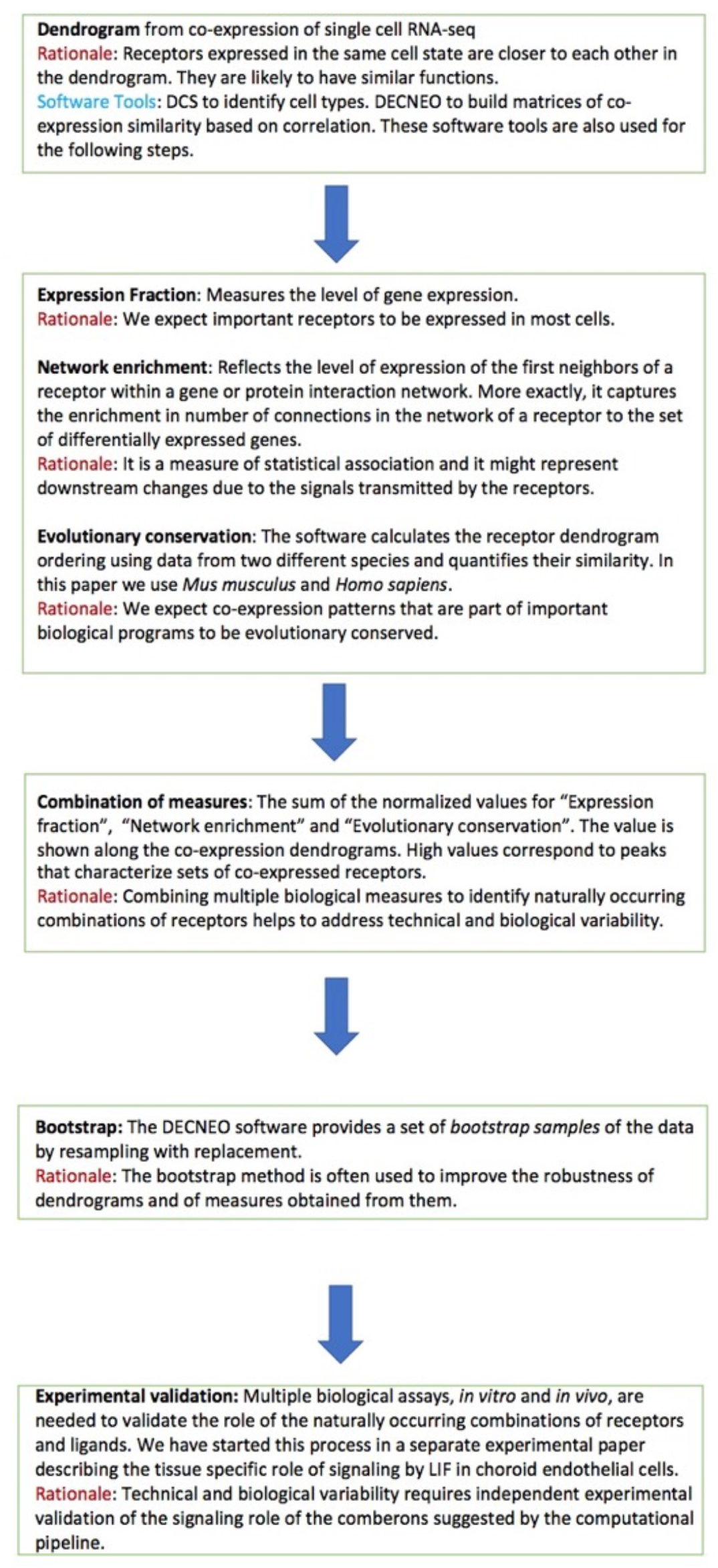
Flowchart with details and rationale of the main steps followed by our process. Additional features not mentioned in the flowchart are:

**Figure 2.**
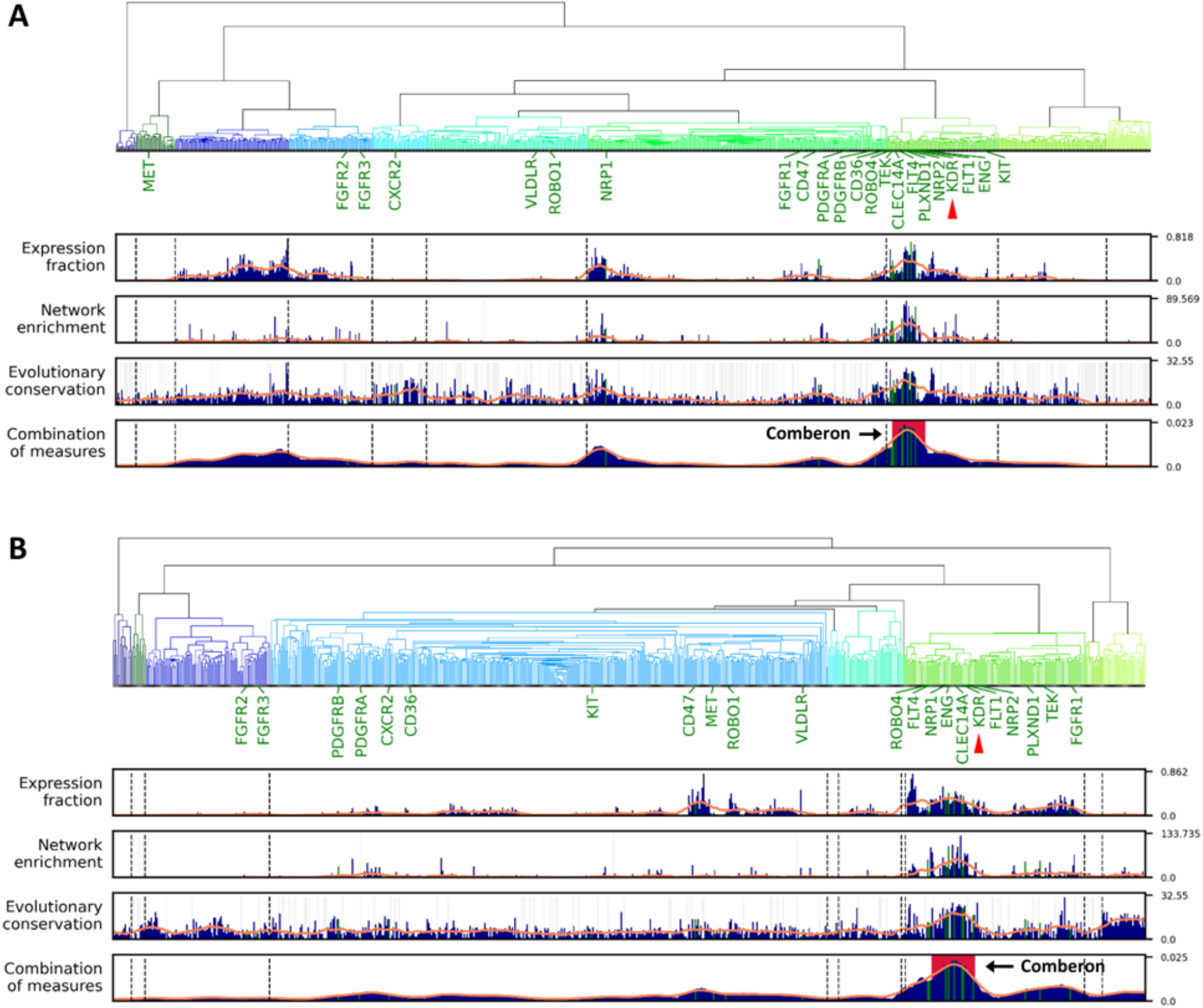
Dendrogram analysis identifies naturally occurring combinations of angiogenesis receptors in endothelial cells. The top of panels A and B displays the dendrogram ordering by single cell co-expression of receptors in endothelial cells for: (A) *Mus musculus* and (B) *Homo sapiens*. Receptors that are co-expressed in individual cells are closer to each other in the dendrogram order. Known angiogenesis receptors are labeled under the dendrogram. The red pointer indicates KDR, the main VEGFA receptor. The panels immediately below the dendrogram show three measures: Expression fraction, Network enrichment and Evolutionary conservation. The lowest panel shows the combination of these three measures. The solid orange line in each panel is a moving window average. The largest peak in the combination of measures panel, representing the main comberon (abbreviation for naturally occurring combination) is shown with a red background.

We also introduce a new software tool, DECNEO (DEndrogram of Co-expression with Evolutionary Conservation and Network Enrichment Ordering), that identifies general and tissue-specific naturally occurring combinations of receptors in endothelial cells and provides visualization and statistical analysis.

## RESULTS

### Overview of the single-cell gene expression data

We used the DCS pipeline [18,19] to process the single-cell RNA-seq PanglaoDB database [20]. The pipeline was applied to each sample independently to normalize the data and identify endothelial and non-endothelial cells. There are much more data from *Mus musculus* studies than *Homo sapiens* resulting in higher statistical power in the analysis of *Mus musculus* tissues.

The independent datasets, which are not part of the PanglaoDB database, are listed in Table S5. In the tissue-specific analysis we use the single cells dataset from Voigt et al. [21] containing *Homo sapiens* choroid data from 18,621 cells in 8 samples. The samples were enriched for endothelial cells, mapped by CellRanger and then normalized and annotated by DCS.

### Overview of the process

A flowchart of the process is shown in Figure 1. Comberon is an abbreviation for naturally occurring combination.

- **Visualization:** As shown in Figure 2, the DECNEO software provides visualizations of the dendrogram and panels that plot the measures described above. It also highlights the location in the dendrogram of the known angiogenesis receptors and allows users to inspect individual measures and to identify cases when they are not concordant.
- **Statistical analysis and validation:** Multiple validations and statistical tests are performed, including a permutation test, validations with independent datasets, the use of a set of receptors with known function (which includes VEGF receptors and other receptors previously shown to play a role in angiogenesis) and interface with other common functional enrichment methods using DAVID and Metascape.
- **Application to intercellular communication:** An extension of the method, shown in Figure 5, can be used to analyze ligands and receptor for a pair of cells. Further evidence for the functional importance of naturally occurring combinations of receptors is obtained by identifying the corresponding naturally occurring combinations of ligands.

### Dendrogram of co-expression ordering

In Figure 2 we show the dendrogram obtained from the analysis of the PanglaoDB dataset. We also show the three measures introduced in the flowchart and their combination. Co-expressed receptors are close to each other in the dendrogram.

The known angiogenesis receptors that are expressed in endothelial cells in the PanglaoDB *Mus musculus* datasets are shown in a panel below the dendrogram in Figure 2A. These known angiogenesis receptors localize in the ordered dendrogram near each other and near KDR and FLT1, the two main VEGF receptors. The Likelihood-Ratio test p-value of a logistic regression for the distance of the known angiogenesis receptors from KDR and FLT1 is less than 7 · 10^−4^. The dendrogram ordering therefore provides information about the role of a receptor in angiogenesis. The non-parametric robust PPMLE was also applied to test the distance of the known angiogenesis receptors from KDR and FLT1 in the dendrogram. For both of these receptors the tests gave a p-value < 5.4 · 10^−6^ confirming the results of the parametric test,

### Genes in the main naturally occurring combination and bootstrap analysis

Using the “Combination of measures” of the PanglaoDB *Mus musculus* data we determined which of the 898 expressed receptors are localized in the largest peak identified by this quantity (corresponding to the peak with a red background labeled as comberon in Figure 2). These 38 genes include 10 of the 22 known angiogenesis receptors, among them KDR and FLT1, the two main VEGF receptors. The probability of such an outcome by chance is less than 2.4 · 10^−9^ by hypergeometric test. This indicates that the largest peak captures receptors that are important in angiogenesis. The statistical significance is higher than that for the distance from KDR and FLT1 calculated in the previous section, showing that the naturally occurring combination is a useful concept for functional elucidation of receptor expression.

We performed a boostrap frequency analysis and a permutation test to quantify the significance of a receptor being included in the main peak (the comberon of Figure 2) in the ordered dendrogram. The bootstrap method [22] is often used to improve the robustness of a dendrogram [23]. Table 1 shows the genes with the largest bootstrap frequency. The bootstrap method performs 100 partial resamples and the frequency indicates how many resamples contain the receptor within the main peak (comberon) shown in Figure 2, with a frequency of 1 corresponding to 100%.

**Table 1.**
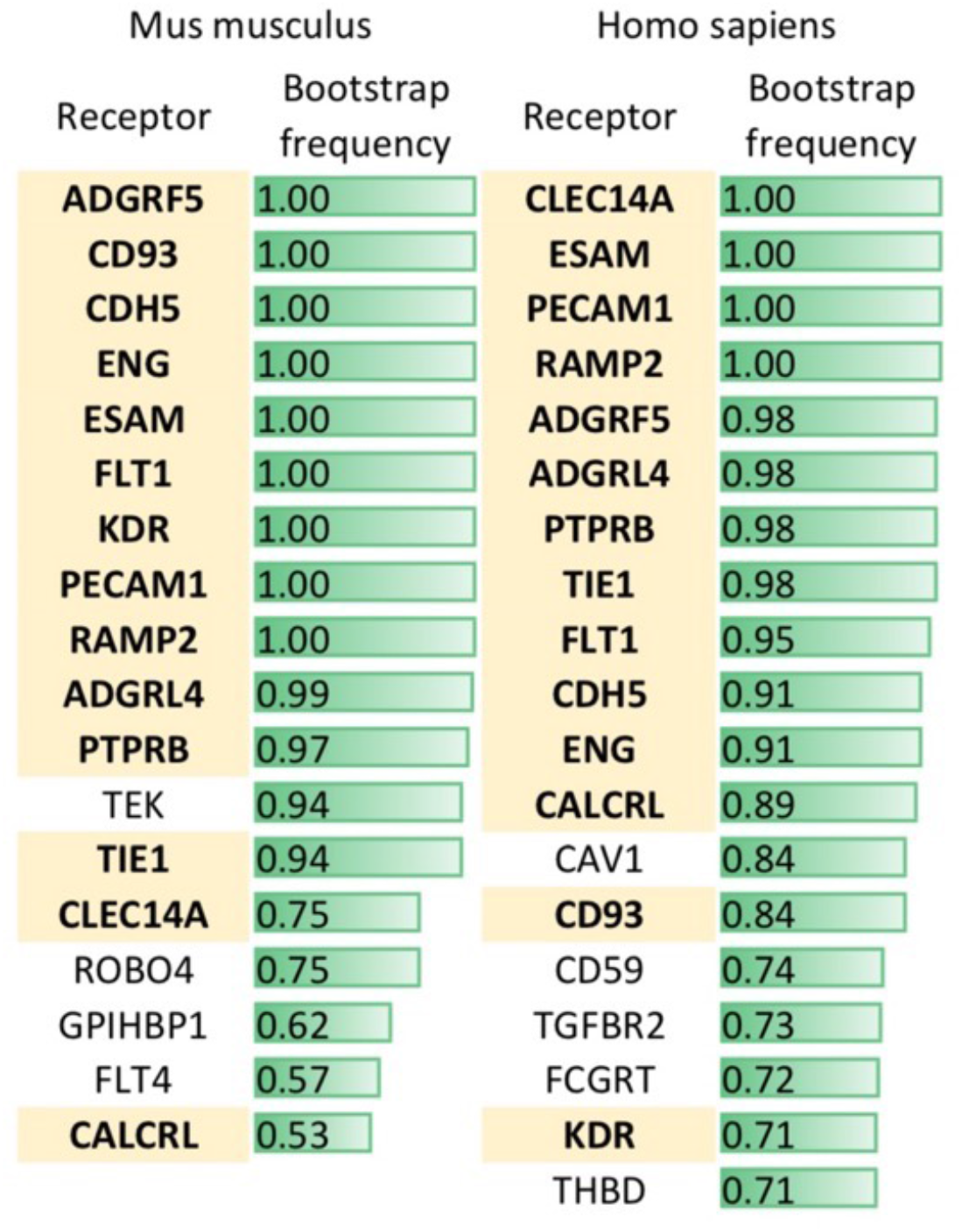
Mouse and human naturally occurring combinations of receptors in endothelial cells are strongly conserved. Main comberon bootstrap frequencies of PanglaoDB endothelial cells annotated by DCS for *Mus musculus* and *Homo Sapiens*. Only receptors with values above 0.5 in mice (this corresponds to a p-value of less than 10^−7^) and 0.7 in humans are shown. Receptors shown in bold with a yellow background are present in both lists and are therefore common to the main comberon of both species.

We used a hypergeometric test for the top genes from the mouse bootstrap shown in Table 1. There are 7 known angiogenesis receptors in this set, which gives a p-value of 5 · 10^−8^. As negative controls we used 30,836 sets of genes from the GO Biological Process class (see Table S3). Table S3 shows that the only strongly significant associations are with terms related to angiogenesis.

Receptors with *Mus musculus* PanglaoDB bootstrap frequencies of at least 0.5 are included in Table 1. A permutation test allowed us to obtain a non-parametric estimate of the probability of obtaining a frequency above 0.5 in *Mus Musculus* as p< 10^−7^. This test shows that the bootstrap method used is extremely unlikely to identify genes with a frequency higher than 0.5 in the absence of co-expression. A lower probability set (p<0.05) can also be identified by including bootstrap frequency values between 0.33 and 0.5. This includes an additional nine genes shown in column 1 of Table S4.

The conservation between mouse and human receptors in the main endothelial comberon, shown in bold with a yellow background in Table 1, was also analyzed with the hypergeometric test and was clearly significant (p< 10^−23^).

### Additional statistical validation with split and independent datasets

Initial statistical validation was obtained by splitting the original dataset into two separate parts and by obtaining two independent datasets. The split was done in two ways, one where the splits had the same number of studies and the other where they had the same number of samples. The results from the split datasets are shown in Table S2. For the splits containing the same number of studies the Pearson correlation of bootstrap frequencies between the two splits was 0.87. The number of genes with bootstrap frequency above 0.5 in the first split dataset that was also found in the second split was 17 out of 23. This gives a p-value of 1.5 · 10^−27^ using the hypergeometric test. For the splits containing the same number of samples the Pearson correlation of bootstrap frequencies between the two splits was 0.91. The number of genes with bootstrap frequency above 0.5 in the first split dataset that was also found in the second split was 16 out of 26. This gives a p-value of 3.3 · 10^−24^ using the hypergeometric test.

As independent datasets for further validation we obtained 16 additional single-cell mice gene expression studies and 6 additional single-cell human studies, not present in PanglaoDB. The bootstrap frequencies for the endothelial cells of the independent dataset were compared to those of the original dataset. The Pearson correlation of bootstrap frequencies between the original and the independent mouse datasets was 0.77 and for the human dataset was 0.78. For human samples, the number of genes with bootstrap frequency in the top 20 in the main dataset that had bootstrap frequency above 0.5 in the independent dataset was 16 out of 20. This gives a p-value of 4.2 · 10^−19^ using the hypergeometric test. For mouse samples, the number of genes with bootstrap frequency in the top 20 in the main dataset that had bootstrap frequency above 0.5 in the independent dataset was 17 out of 20. This gives a p-value of 7.0 · 10^−23^ using the hypergeometric test.

### Tissue-specific receptors

There are eight *Mus musculus* and *Homo sapiens* tissues within the PanglaoDB database annotated by DCS with substantial data (at least two studies with at least 300 endothelial cells each including at least five samples). The correlation among the 21 studies within these eight tissues is shown in Figure 3. Most studies belonging to the same tissue cluster together. Additionally, the hypothalamus, olfactory bulb and subventricular zone studies, which are all parts of the brain, also cluster together. The average correlation values within tissues of the same type are almost always larger than the average correlation with other tissues (see Table S9).

**Figure 3.**
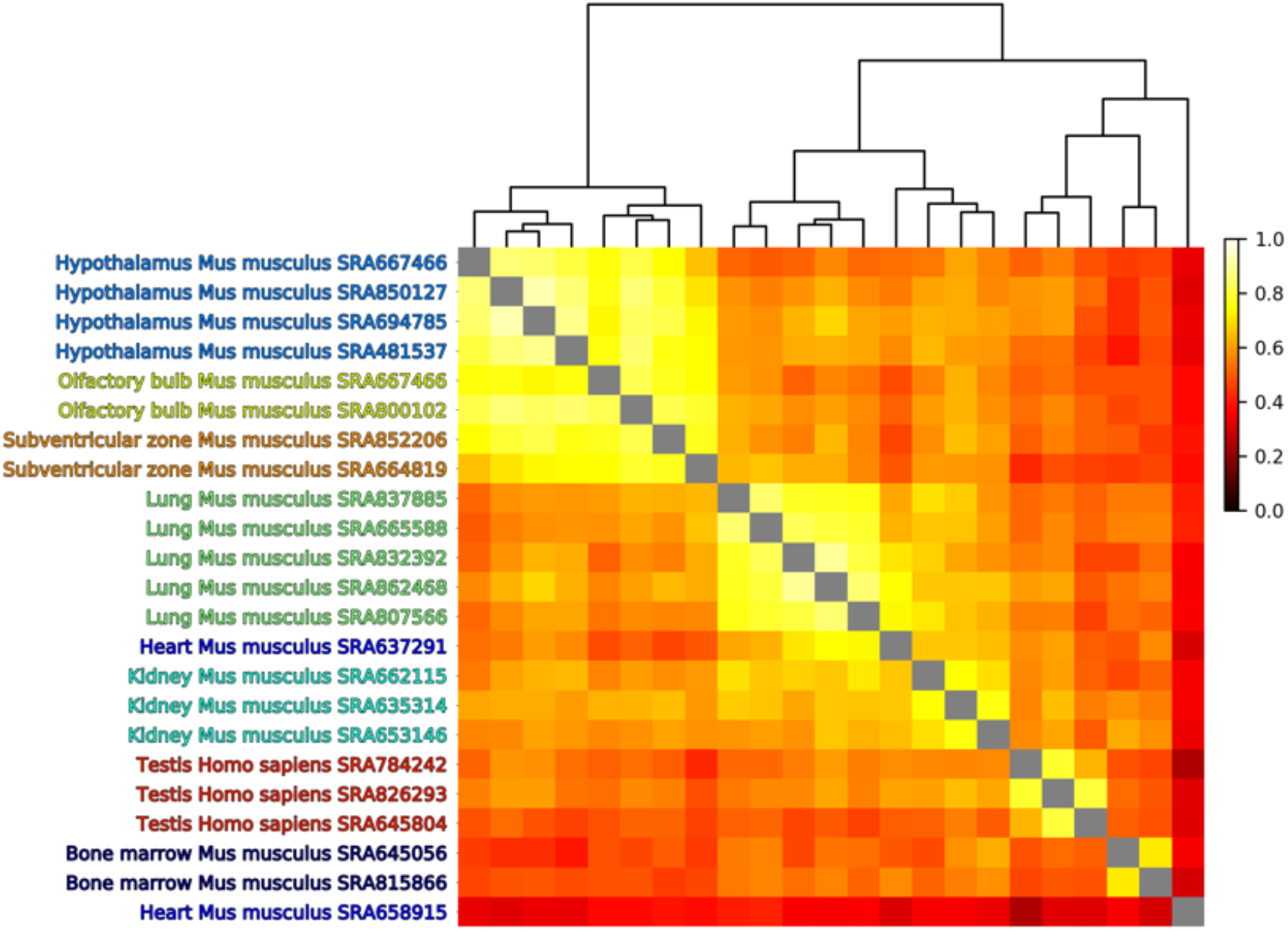
Analysis of endothelial cells from selected studies in PanglaoDB database annotated by DCS. Clustering of Pearson correlation of the largest peak bootstrap frequencies for the combination of measures. Studies from almost all similar tissues group together.

Figure 4 shows the receptors with the highest bootstrap frequencies (included in the comberons) for the seven tissues shown in Figure 3 and for one additional human dataset obtained from the choroid of the eye [21]. These tissues usually also have high bootstrap frequencies for the top 13 receptors from the general analysis of Table 1 (receptors left of the red line in Figure 4). The results shown in Figures 3 and 4 indicate that endothelial cells have partially tissue-specific co-expressed receptors within the comberons.

**Figure 4.**
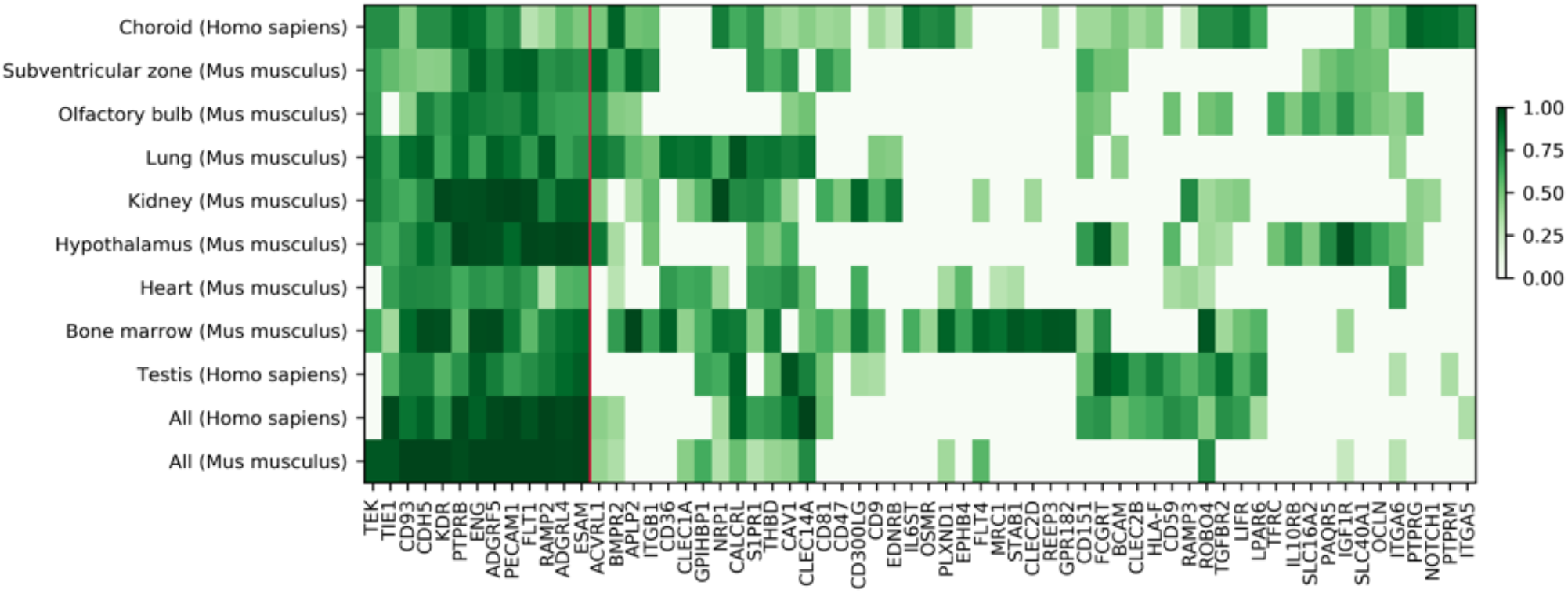
Comberons of endothelial cells of specific tissues from PanglaoDB and choroid. Under All we include the global analysis shown in Table 1 for comparison. The 13 receptors to the left of the red line are the 13 top receptors from Table 1 (with a boostrap frequency higher than 0.9). The rest of the receptors are within the top-10 tissue-specific genes for at least one set.

### Choroid endothelial receptors

Given the heterogeneity between tissues and the presence of partially tissue-specific receptors in the comberons we analyzed in depth the main comberon of human choroid endothelial cells, also in the light of very recent experimental results we have obtained [17] and of the therapeutic relevance. The choroid vessels of the eye play an important role in the development of Age-related Macular Degeneration (AMD) one of the leading causes of blindness [17]. Several VEGF inhibitors are used as FDA-approved therapies for AMD but not all forms of the disease benefit from them [17].

Table 2 shows the 25 receptors with a bootstrap frequency higher than 0.7 in human endothelial cells (see also figure S14). Single cell RNAseq data were from Voigt et al [21]. Out of these 25 receptors, 6 have the strongest evidence of relevance to choroid vasculature, either because they are targets of approved AMD drugs (KDR and its co-receptor NRP1), are associated with increased AMD risk in GWAS studies (CFH) [24], or have been shown to regulate choroid angiogenesis in our separate experimental paper [17] (LIFR, IL6ST). Out of the remaining 19 receptors, 16 have at least ten publications associating them with angiogenesis in a systematic Pubmed search. KDR, the main VEGF receptor, is by far the most studied. While the scientific community has accumulated substantial knowledge about the role of multiple receptors in angiogenesis, this knowledge is incomplete. We did not find published datasets examining them under the same experimental conditions and in combinations of multiple sizes.

**Table 2.** The receptors in the main comberon of human choroid endothelial cells. All the receptors with substantial evidence for a role in angiogenesis are shown in red.

| Receptors | Frequency | Pubmed angiogenesis | Strongest Experimental evidence |
| --- | --- | --- | --- |
| **BMPR2** | **0.9** | **92** |  |
| PTPRG | 0.9 | 3 |  |
| PTPRB | 0.89 | 9 |  |
| **NOTCH1** | **0.87** | **388** |  |
| PTPRM | 0.86 | 0 |  |
| **ENG** | **0.85** | **891** |  |
| **IL6ST** | **0.82** | **13** | **LIFR co-receptor** |
| **LIFR** | **0.82** | **8** | **Our LIF experimental paper (Li et al., 2021)** |
| **NRP1** | **0.8** | **399** | **KDR co-receptor** |
| **PLXND1** | **0.78** | **31** |  |
| **ITGA5** | **0.77** | **18** |  |
| **OSMR** | **0.75** | **9** |  |
| **PECAM1** | **0.75** | **404** |  |
| **CD46** | **0.74** | **22** |  |
| **CDH5** | **0.74** | **51** |  |
| **KDR** | **0.74** | **1952** | **main VEGFA receptor** |
| **ROBO4** | **0.74** | **74** |  |
| **TEK** | **0.74** | **210** |  |
| **TGFBR2** | **0.74** | **194** |  |
| **TIE1** | **0.74** | **205** |  |
| **ADIPOR2** | **0.73** | **11** |  |
| **S1PR1** | **0.71** | **68** |  |
| **CFH** | **0.7** | **69** | **Main genetic risk factor for AMD** |
| **IL4R** | **0.7** | **13** |  |
| **ITGAV** | **0.7** | **26** |  |

### Ligand-receptor pairs in choroid fibroblasts and endothelial cells

Ligands can originate from multiple cell types but finding a combination of ligands within a cell type that at least partially correspond to the combinations of receptors we have identified in endothelial cells would support the validity of our approach to intercellular communication. This is indeed shown for choroid fibroblast ligands and for choroid endothelial receptors in Figure 5.

**Figure 5.**
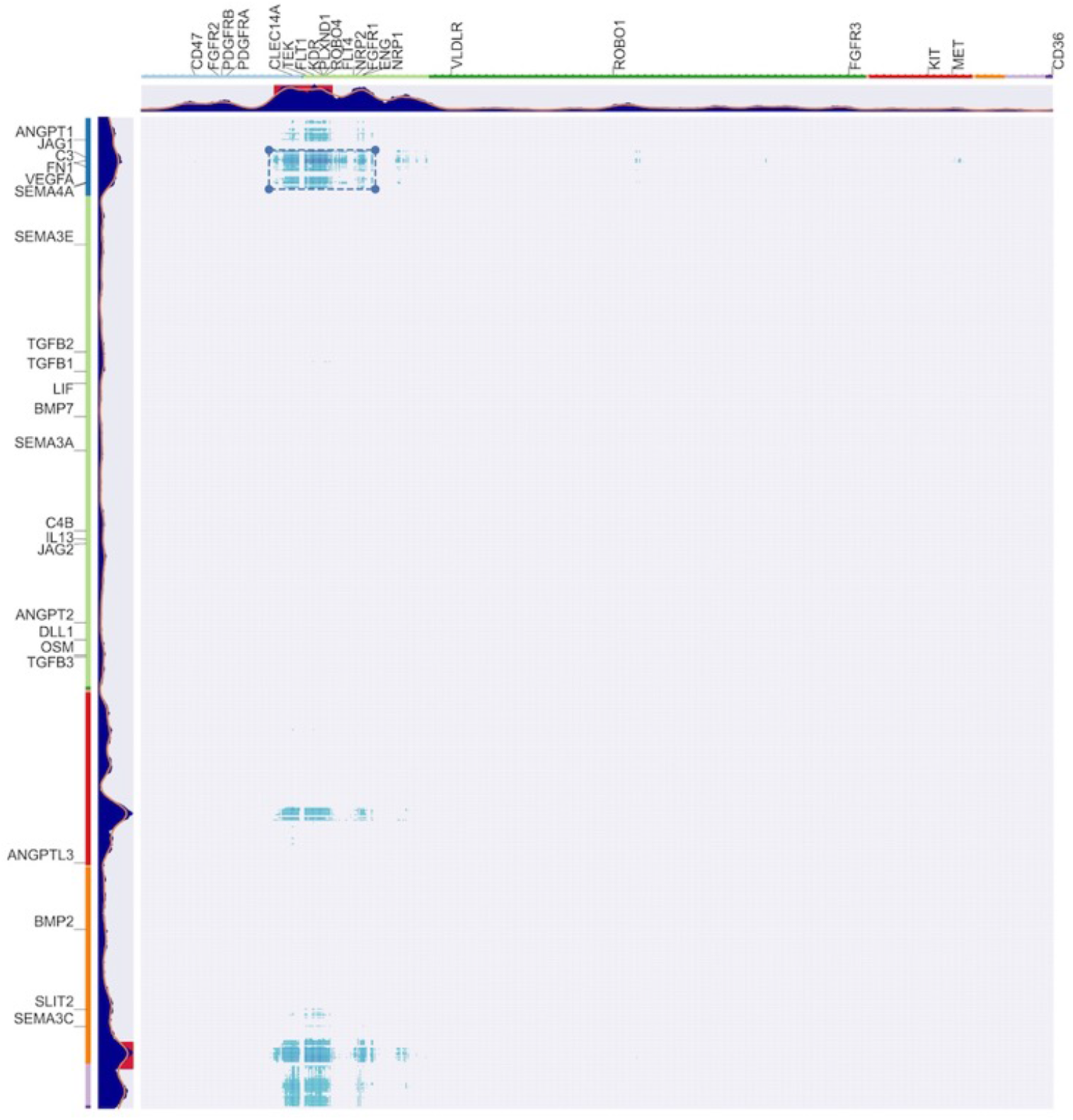
Two-dimensional matrix of receptor co-expression in human choroid endothelial cells and of the corresponding ligands co-expression in human choroid fibroblasts. Ligands or receptors co-expressed in individual cells are closer to each other in the dendrogram order. The peaks represent the combination of measures, as in the dendrograms of Figure 2. Endothelial receptors are on the x-axis and fibroblasts ligands on the y-axis. Known choroid endothelial receptors labels are shown on the horizontal axis and the corresponding ligands are shown on the vertical axis. The box indicates the region with the highest combinatorial communication measures.

Furthermore, if two cells communicate using combinations of signals, ligand-receptor pairs between a comberon of ligands in one cell type and a comberon of receptors in the other cell type are expected to be enriched in “known ligand-receptor pairs”. These “known ligand-receptor pairs” are supported independently, for example by protein-protein interactions datasets. One of the most widely used lists of “known ligand-receptor pairs” is that of Ramilowski et al [25,26] and we have also used this list in our analysis.

The first result supporting our hypothesis is the strongly significant correlation between the “combination of measures” value for a ligand-receptor pairs (obtained averaging the ligand and receptor values) and the enrichment in “known ligand-receptor pairs” in its dendrogram neighbors. For the datasets of Figure 5 the p-value for both Pearson (correlation coefficient 0.4) and Spearman (correlation coefficient 0.33) was < 2.2 · 10^−16^.

The analysis shown in Figure 5 identifies a region where the mean of these quantities (the “combination of measures” for a ligand and a receptor and the enrichment in “known ligand-receptor pairs” in its dendrogram neighbors) is maximal. This region corresponds to the main comberon of the endothelial receptors and to one of the comberons of the fibroblast ligands. This finding is consistent with the fact that angiogenesis signaling is only one of several possible roles for fibroblasts. The ligand and receptors within this region are enriched for genes with known function in angiogenesis (GO Biological Process, GO:0001525 angiogenesis, FDR 1.4 · 10^−8^).

In Figure 5 we also show labels for known choroid endothelial angiogenesis receptors and for the corresponding ligands. Note that VEGFA and its ligand KDR are included in the box with the highest combinatorial communication measures.

## DISCUSSION

The study of correlation in gene expression data is a well-known and fruitful approach that has contributed to the understanding of gene function, for example using software methods like WGCNA (weighted gene co-expression network analysis) [27]. The DECNEO method we suggest here has been designed explicitly to decode intercellular communication using single-cell RNAseq data. The main differences with WGCNA can be appreciated by comparing the flowchart in Figure 1 of the initial WGCNA paper [27] with the flowchart shown in Figure 1 of this paper. One of the main differences is the use of additional biological information to identify the sets of genes in DECNEO rather than just to characterize them as in WGCNA. Especially relevant to the study of receptors is the enrichment of connections to up-regulated genes (network enrichment), which might depend on the downstream changes due to the signals transmitted by the receptors. In general, the method deals with the noise of biological data by averaging information from different features that we expect to be associated with combinations of intercellular signals. As it can be appreciated from Figure 2, these features are layered on the main framework given by the ordering by co-expression within the dendrogram. This Figure also shows the visual output of the software which assists in the exploration of the data. The addition of bootstrap analysis is a useful statistical feature based on the experience of many scientific fields where dendrograms are used [23].

The strong conservation between species shown in Table 1 and in the related statistical analysis suggests that the combination of receptors has a functional role which natural selection has been under pressure to preserve. The significant enrichment for many receptors with a known function in angiogenesis indicates what this role is likely to be.

While anti-VEGF therapies have had a major impact, the use of VEGF to promote angiogenesis and endothelial survival has not been clinically successful [6,5,4]. This is partly due to the fact that the vessel growth stimulated by VEGF alone is abnormal and accompanied by leakage [5,4,28]. These observations also suggest that VEGF is only part of the signal for normal vessel growth and that a broader combination of receptors is needed to transmit the signal of normal angiogenesis.

As we mentioned in the Introduction there is growing consensus that drug combinations acting on multiple targets are also needed to improve clinical response to angiogenesis inhibitors [1,4,5]. Among the strategies at a more advanced stage is the development of Facirimab, a dual inhibitor of VEGF and of the Ang/TEK pathway [29]. Note that the TEK alias name TIE2 is often used in the literature [29]. Promising preclinical and clinical trial data in age-related macular degeneration have been obtained [29] even if in clinical trials Facirimab is not always clearly superior to VEGF inhibition alone [30]. Table 2 does indeed list TEK as part of the main naturally occurring combination of receptors from human choroid endothelial cells.

The last part of the method consists of a series of in vitro and in vivo experimental tests exemplified by a study of the action of LIF on choroid endothelial cells described in our experimental paper [17]. The LIF receptor LIFR is also listed in Table 2, again providing more support for the relevance of the list, as in the case of TEK.

Table 2 also shows that a substantial literature has accumulated on the role in angiogenesis of many of the receptors listed there. We found this knowledge to be accumulating in a fragmentary manner, mainly consisting of studies of individual receptors and, to a lesser extenct, of small combinations. This bias in our knowledge can be explained by the size of experiments that is reasonable to accomplish within an academic lab. It could be addressed either by a dedicated larger effort or by an increased appreciation by the scientific community of the importance of partial steps towards solving the complex problem of combinatorial intercellular communication.

The process used by Shinya Yamanaka and his group to identify combinations of factors to induce pluripotent stem cells can be used as an informative analogy for the importance of decoding combinatorial cell to cell signaling [15]. Starting from a longer list of transcription factors Yamanaka used several methods, including a computational analysis of gene expression, to identify a set of 24 transcription factors that he then reduced experimentally, finding that just 4 of them were sufficient [15]. Signaling by transcription factors, however, can induce switches between cell types (which are not common in differentiated cells) while receptors regulate changes between cell states, for example from quiescence to proliferation. Yamanaka has shown that signaling through transcription factors is characterized by multiple signals that need all to be present for an effector function. In the case of receptors, we know that some of them (like the VEGF receptors) can induce a strong response on their own, but that integration of multiple signals does also take place. While different strategies might be needed in the case of receptors, it is still reasonable to expect that a computational analysis of expression data, followed by a systematic experimental search, might lead to the decoding of intercellular combinations of signals.

### Possible limitations

We believe the results reported here to be potentially useful but we also would like to point out cases where they should not be overinterpreted. The combinations of receptors we have reported should not be used to exclude the importance of a receptor for angiogenesis. This is not a necessary inference from our results, because cells might communicate using individual ligands or receptors in addition to using combinations. Furthermore, we are far from having investigated every possible tissue or condition.

It is also not justified to conclude that a receptor that is part of the naturally occurring combinations we have identified should have an important role in angiogenesis when engaged in isolation. While this might in some instances be the case, only appropriate experimental tests will be able to clarify the role of a receptor when stimulated together with the other members of the combination. It is certainly possible that the magnitude or even the direction of the effects on endothelial cells might be modified by the presence of other signals.

In addition to publication this project might benefit from adopting a format suitable for continuous updating, to keep up with the increasing amount of available single cell data. These data are likely to keep growing at even faster rate, considering that the identification of naturally occurring combinations can be extended to other cell types, go beyond receptors (with some modifications in the methods) and include more species to measure conservation.

We have only investigated the more general properties of receptor combinations in endothelial cells. For example, some receptors might belong to more than one combination. Our results on tissue-specific receptor analysis provide the first step in this direction, but there might be combinations associated with different cell states within a tissue, including disease states. More data will certainly improve the precise identification of the receptors included in the combinations, especially at the organ and tissue-specific level. It will also be interesting to use the tools we describe to study how aging affects intercellular communications. Recent data show, for example, that angiogenic signals from fibroblasts to endothelial cells change with aging [31] and VEGF signaling has been shown to affect the aging process [32,33]. Subtypes of endothelial cells could also be studied.

It is important to promote scientific replication, not only internally as we have often done, but also externally. We facilitate this by sharing our software code in an organized format.

### Conclusions

This paper identifies naturally occurring and evolutionarily conserved combinations of receptors in endothelial cells that are likely to play a role in angiogenesis and describes a software method applicable to combinations of receptors in general. We also show tissue-specific combinations and, in the case of choroid endothelial cells, consistency with well-established and recent experimental findings, some of which are reported by us in a related experimental paper [17].

## MATERIALS and METHODS

### Data sources

Three sources of data were used in our analysis:

1. The primary data source in our study was PanglaoDB [20], a database of scRNA-seq datasets of *Homo sapiens* and *Mus musculus* studies, that collects 22,382,464 cells from 236 studies (SRAs, or Sequence Read Archive submission accessions) in 1368 samples (SRSs, or Sequence Read Archive sample accessions), spanning numerous tissues and cell types (see Table S5). After quality control PanglaoDB provides data from 4,459,768 *Mus musculus* cells and 1,126,580 *Homo sapiens* cells. Each of the datasets was downloaded from NCBI SRA and mapped to the GRCm38/GRCh38 genome assembly by the authors of PanglaoDB [20]. All cells in PanglaoDB are annotated with cell type labels by Alona [34], a web-based bioinformatics tool for scRNA-seq analysis. This cell type annotation is incorporated in the PanglaoDB database. In addition to the Alona annotation, we processed each of the datasets in PanglaoDB using a platform for cell-type annotation that we have recently developed [18,19], DCS 1.3.6.10.
2. We also obtained an independent dataset for validation, composed of 16 single cell mice gene expression studies and 6 single cell human gene expression studies (see Table S5). This independent dataset was obtained after the analysis of the main dataset had already been completed and did not affect any of the methodological choices.
3. Another source of data was a dataset from Voigt et al. [21] containing *Homo sapiens* choroid scRNA-seq on 18621 cells in 8 samples enriched for endothelial cells. The dataset was downloaded from NCBI and re-mapped with CellRanger 4.0.0 and the GRCh38 genome assembly; each of the 8 samples has 4 paired-end reads sequencing runs (SRRs) merged into one. In our alternative implementation of DECNEO, used as a validation, we used the original Voigt et al. [21] data mapping, quality control, and cell-type annotation of the choroid cells.

### Input data format

See Figure 6 for a graphical summary of the analysis pipeline. The method is designed to take two single-cell transcriptomics datasets corresponding to two different species, *Mus musculus* and *Homo sapiens*. Each input datasets must be in a table format where gene identifiers are in rows and cells and samples identifiers are in columns. Where possible, gene identifiers should be converted to one species, e.g. *Homo sapiens*, using their homologs. Starting from the gene counts matrix, the gene expression values should be scaled so that the sum of values of each cell in the dataset is identical (we recommend a scaling factor of 10^5^) and then log-transformed. Cell type and sample annotation are required for each cell in each dataset, assigning cells to two different classes, e.g. endothelial and non-endothelial cells. The preprocessing step described above is necessary for the analysis pipeline. The preprocessing step is not directly included in DECNEO, but DCS [18,19], can be used to normalize and annotate cells.

**Figure 6.**
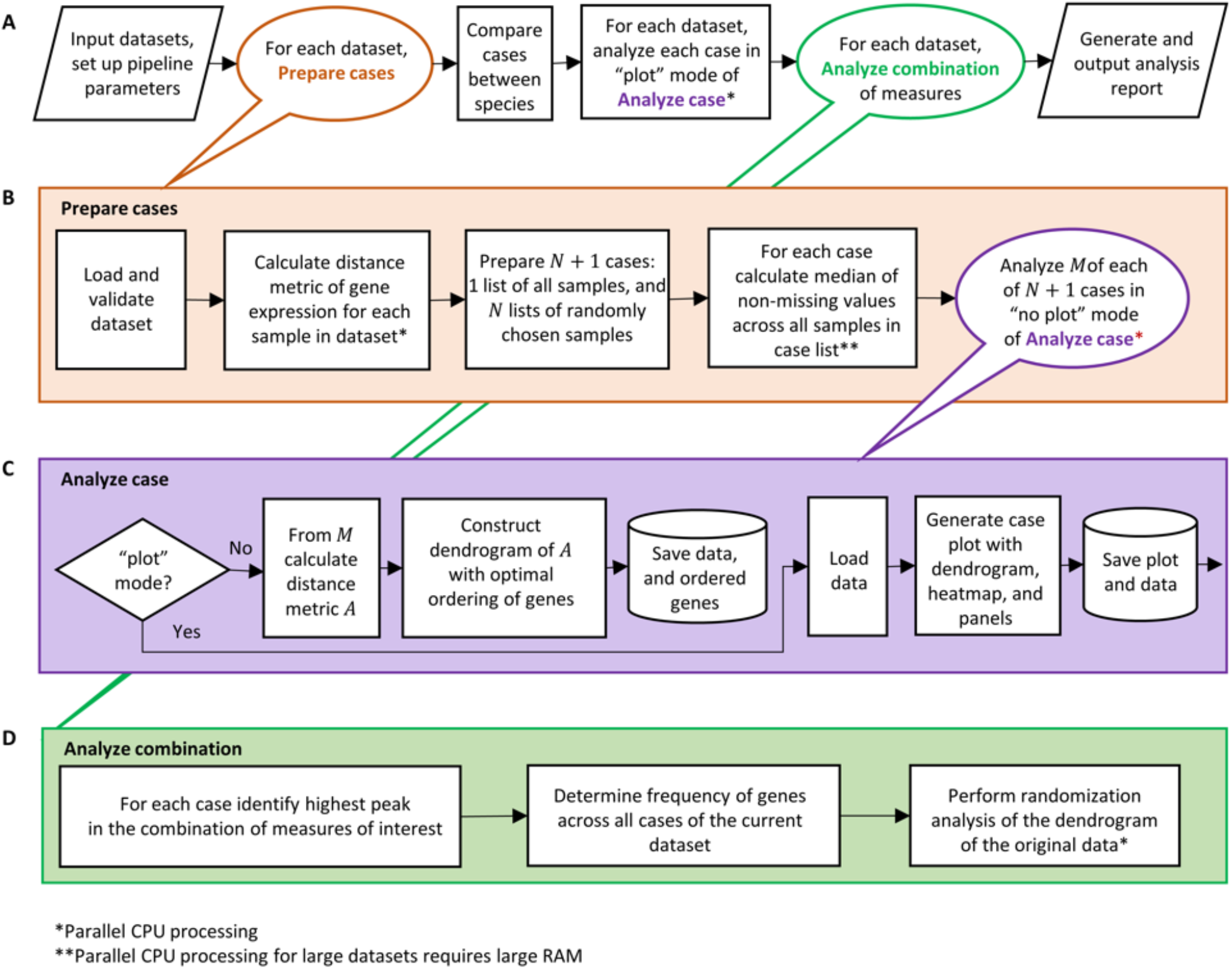
Summary of the analysis pipeline. Normalized single cell transcriptomics datasets are processed in steps depicted in panel A. The principal steps of the pipeline are outlined in panels B, C and D. An analysis report is generated at the end of the pipeline.

### Dataset validation and bootstrap resamples preparation

First, the dataset is validated and checked for the inclusion of a sufficient number of samples (see Figure 6). If the number of samples in a dataset is below five we combine them and then split the data into ten non-overlapping sets, to allow bootstrapping. Next, within each sample = 1 … *k*, we calculate a matrix 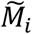 of distances between gene pairs based on gene expression according to a user-specified metric (Euclidean, cosine, or correlation distance). In a given sample, only genes expressed in at least 5% of cells are considered. For the choroid dataset the cutoff is 1% because of the high quality of this dataset, which was experimentally enriched for endothelial cells and had a higher sequencing depth (22). For each sample, the distance is calculated between each gene in a set of *m* genes of interest (in Figure 2 the genes of interest are all known receptors) and each gene in the complete set of *n* expressed genes, resulting in *M*, an *n* × *m* matrix that contains the *n* genes in rows and *m* receptors in columns. Samples (i.e., the set of *k* matrices, one per sample) are then aggregated using the median distance among non-missing values. For missing values in the distance matrix *M* we assume no correlation or, in the Euclidean case, replace with the maximum of *M*. When the matrix *M* is calculated using all the available samples, we refer to this as the original data. In addition to the *original data*, we prepare a set of *bootstrap resamples* by resampling with replacement all samples *N* times (we used *N* = 100): i.e. we randomly choose with replacement *k* samples and define a distance matrix following the same procedure as in the *original data*. The size of the bootstrap resamples was validated for one dataset, the choroid dataset, where we compared the results of 100 and 1000 resamples. The output of the bootstrap was very similar among the two independent resamples. The bootstrap frequency Pearson correlation between them was 0.99. The set of bootstrapped distance matrices is used to analyze the effect of gene expression variability and for statistical inference, as explained later in the statistical analysis section.

### Dendrogram ordering by co-expression

The ordering of the dendrogram (see Figure 2) is based on the Pearson correlation of normalized single-cell gene expression, which is a commonly used gene co-expression measure. The similarity metric is the Euclidian distance of the Pearson correlation. We also compared alternative choices for these metrics. For example we tested Spearman and Pearson correlations of gene expression (see Table S1).

Here, an *m* × *m* symmetric matrix *A* with no missing values is calculated from each *n* × *m* matrix *M* by considering the Euclidean distances of the Pearson correlations between the columns of *M*, thus defining a distance matrix for the *m* receptors only. We then perform an agglomerative clustering on the matrix *A* using a Ward variance minimization linkage method [35] and construct a dendrogram (alternative clustering similarity metrics and linkage methods are compared in Table S1). Correlation is compared to the Euclidean distance of Pearson correlation in the dendrograms shown in Figures S1, S2 and S3. Finally, receptors, which are the leaves of the dendrogram, are ordered to minimize the sum of the distances between adjacent leaves in the dendrogram layout (25). This dimensionality reduction procedure defines a “dendrogram distance” between any two ordered receptors, which is a positive integer equal to the difference of their ordering index. The *N* bootstrapping resamples and the original data are processed according to the steps shown in Figure 6.

Following this procedure, genes co-expressed in individual cells are closer to each other in the dendrogram order.

### The 3 measures shown in the lower half of the Dendrogram Figure (Figure 2) are

#### 1 Expression fraction

For each receptor we calculate the fraction of cells that express that receptor at a non-zero level. We calculate the median across the samples. We expect important receptors to be expressed in most cells.

#### 2 Network enrichment

To find receptors close to genes that are differentially expressed in endothelial cells we use a predetermined undirected gene interaction network known as *Parsimonious Composite Network* (PCN) [36]. This network contains 19,783 genes and was obtained by combining data from 21 popular human gene interactions networks. Only interactions supported by a minimum of two networks were included. The 21 networks included interactions derived from multiple methods, such as protein-protein interactions, transcriptional regulatory interactions, genetic interactions, co-expression correlations and signaling relations. The authors extensively evaluated the PCN network showing that it was superior to any of the 21 networks used to build it. We independently determined that sets of genes with similar functions are close to each other within this network. Using inflammation, cell cycle and metabolic gene sets we measured intraset efficiency, a metric we have previously introduced [10], and the enrichment for direct connections between the genes in a given set using the binomial distribution. All the tests showed highly significant p-values, supporting the high quality of the PCN network.

First, we determine the set of the top 1000 differentially expressed genes, *S*, in a dataset consisting of several samples. The differential gene expression for each sample is computed using a t-test between the gene expression of endothelial cells and non-endothelial cells in that sample. Genes with a p-value below 0.001 are retained and ordered according to the decreasing t-test statistic. The median of the ranks across samples is taken to rank the differentially expressed genes of the entire dataset.

The t-test was also compared with the Mann-Whitney U test and the network enrichment obtained was very similar (see Table S1 for bootstrap frequency obtained with the two methods).

To obtain a p-value we use the binomial distribution. Let *p* be the probability that a randomly chosen PCN edge is connected to a gene in the set *S* of differentially expressed genes. We estimate *p* as the number of edges with at least one gene in *S* divided by the total number of edges in the network. For each receptor *j* the probability of having *k*_*j*_ or more enriched edges out of its total degree *n*_*j*_ in PCN is given by

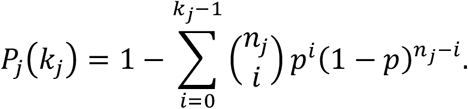

The negative logarithm in base ten of this probability is our method’s “Network enrichment” measure.

This measure reflects the level of expression of the first neighbors of a receptor within the PCN network. More precisely, it captures the enrichment in the number of connections of a receptor to the set of differentially expressed genes.

#### 3 Evolutionary conservation

To evaluate evolutionary conservation, we calculate the receptor dendrogram ordering using data from two different species and we quantify their similarity. The comparison is made by pairing data from different species. In this paper we use *Mus musculus* and *Homo sapiens*. For each receptor *j* in the first case (for example *Mus musculus*) we count the number of receptors in a particular window in the dendrogram ordering (here of size 50, centered at *j*) that are also present in a window of the same size centered at homologous receptor *j* for the second case (for example *Homo sapiens*). The number of overlapping receptors in the neighborhoods of the receptor *j* defines a local measure of evolutionary conservation. Genes that do not appear in both species are marked as missing with a grey bar.

### Other measures

We have considered other measures, including evolutionary rate and age, the average gene expression, the distance between neighboring leaves along the dendrogram, evolutionary conservation of known angiogenesis receptors, and known angiogenesis receptors from gene ontology angiogenesis terms. These measures did not help in identifying the comberon or in the enrichment for known angiogenesis receptors. The main implementation of DECNEO allows for including these measures, or even new, user-defined, measures in future analysis.

### Known angiogenesis receptors

We use for validation a set of 22 receptors that are known to regulate angiogenesis: CD36, CD47, CLEC14A, CXCR2, ENG, FGFR1, FGFR2, FGFR3, FLT1, FLT4, KDR, KIT, MET, NRP1, NRP2, PDGFRA, PDGFRB, PLXND1, ROBO1, ROBO4, TEK and VLDLR. These receptors were obtained before the analysis from a systematic Pubmed search for receptors with functional effects on angiogenesis. The list includes FLT1 and KDR, the main VEGF receptors, and also other targets for anti-angiogenic drugs under development [1,5,2,37]. As negative controls we used 30,836 sets of genes from the GO Biological Process class (see Table S3).

### Combination of measures and comberons

Figure 2 shows how we analyze the combinations of different measures. The lowest panel in Figure 2 shows a linear combination of the measures. “Combination of measures” is a sum of “Expression fraction”, “Evolutionary conservation”, and “Network enrichment”.

For each dataset *d* and for each resample *c*, we first normalize these measures so that the sum of each measure is one and take a moving average along the dendrogram to smooth the normalized measures (using a centered window of 21 receptors and a uniform averaging kernel). Then, we add their values and take a moving average of the combined measure to obtain a combination score 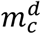.

In the analysis, we identify the highest peak *p*_0_ in 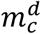, such that *p*_0_ is 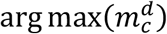, along the dendrogram and we find all the receptors with 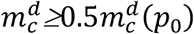 that are in the immediate proximity of this main peak. Then, we count how many times a given receptor is found in the main peak across all bootstrap resamples in a dataset and we calculate its frequency of peak inclusion. The software can also calculate the frequency of inclusion for the main and all other secondary peaks, *p*_*i*_ where *i* ≥ 0, in the dendrogram by considering all extrema of height and prominence larger than 0.05 separated by at least 50 receptors. The prominence of a peak is defined as the vertical distance between the peak and the lowest contour line encircling it but containing no other higher peaks. When secondary peaks are used, each peak *i* (main or secondary) contains at most 50 receptors that are in the immediate proximity of the peak and have 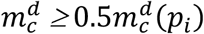, see Figures 2 and S4.

The sets of genes that are included in the peaks are the comberons. It is important to observe that the measures used to identify the peaks are not necessarily increased for a particular gene. For example, as shown in Figure 2, a gene with a low value of network enrichment can be included in a peak because it is co-expressed with other genes that have a high value for this measure.

### Visualization

A “plot” mode (see Figures S1 and 2) can be selected to visualize dendrograms, heatmaps of the *A* matrix (as an option), and several other panels. These panels plot the measures described above, such as the Evolutionary Conservation and Expression Fraction, smoothed using a moving average along the dendrogram (indicated by an orange line), the combinations of these measures and highlight the location in the dendrogram of the known angiogenesis receptors.

### Statistical analysis of Dendrogram Ordering

The likelihood-ratio test was used to analyze the distance in the dendrogram of known angiogenesis receptors from KDR and FLT1. The null hypothesis here is that the known angiogenesis receptors are uniformly spread in the optimally ordered dendrogram. In contrast, the alternative hypothesis is that the known angiogenesis receptors should be in the vicinity of KDR (or FLT1) in the dendrogram, showing that known angiogenesis receptors are co-expressed. Under the null hypothesis every dendrogram receptor has an equal probability of being one of the known angiogenesis receptors. With the alternative hypothesis we use a logistic regression to get the probabilities of the receptors to be a known angiogenesis receptor given the optimal dendrogram ordering. The p-value is calculated using the survival function of a chi-squared continuous random variable.

An alternative non-parametric method was also used for the same purpose [38]. The robust Poisson pseudo maximum likelihood estimation (PPMLE) was implemented in R using the package *gravity* [39]. The p-value was obtained from the difference of the deviances of the null hypothesis and alternative hypothesis using the chi-square distribution.

### Statistical analysis of Comberons

The main aim of this statistical analysis was to show that there is a set of receptors significantly associated with the largest peak of Figure 2. We call the set of receptors associated with the largest peak the main comberon.

The first step was the use of the bootstrap method [22] to improve the robustness of the dendrogram. This is a common procedure in many scientific fields where dendrograms are used [23]. In fact, the paper describing the application of the bootstrap method to dendrogram analysis is one of the 100 most highly cited papers of all time [40,41].

The bootstrapping involved 100 resamples with replacement. The values shown in Table 1 are bootstrap frequencies, where 1 means that the receptor was found in the largest comberon in all of the 100 resamples. See Table S4 for the complete list of the bootstrap frequencies.

We then performed several statistical tests:

1. A permutation test (also called randomization test) [42,43] for the co-expression that was used to build the dendrogram. We wanted to determine the statistical significance of the frequencies in the first column of Table 1, which contains the *Mus musculus* data analyzed with DCS, and we therefore used a control where the ordering of the dendrogram was no longer dependent on the correlation of the receptors’ gene expression. First, we determined the maximum frequency of genes included in the main peak for 100 instances (to match the bootstrapping procedure) with randomized ordering of the dendrogram shown in Figure 2A. We then repeated this procedure for 10^7^. realizations. This calculation was done using the High-Performance Computing Center (HPCC) supercomputer at Michigan State University. From the 10^7^. realizations we obtained a distribution of the maxima of the bootstrap frequencies for the original data case. We did not observe a frequency above 0.5 in any of the 10^7^. realizations. The maximum observed frequency (0.49) was slightly below this number as shown in Figure S5. This procedure allowed us to obtain a non-parametric estimate of the probability of getting a frequency above 0.5. Note that this is a conservative estimate of the p-value given, because it is an estimate for a single value being larger than 0.5, while in the first column of Table 1 there are 18 genes above this threshold. We also empirically determined that this distribution resembles a generalized extreme value distribution, which is a known distribution of maxima (see Figure S5).
2. As a validation we split the dataset into two equal parts. This was done twice, once splitting the samples and once splitting the studies. The bootstrap frequencies of the two splits were then compared using the hypergeometric test.
3. The significance of enrichment of the comberon for known angiogenesis receptors was calculated using the hypergeometric test. This was done for both the original and the bootstrap frequency of at least 0.5.
4. We also obtained an independent dataset composed of 14 single-cell mice gene expression studies, containing 71 samples and 65,057 endothelial cells (see Table S5). The endothelial cells were identified using DCS as was done for the original Panglao dataset. This independent dataset was obtained after the analysis of the main dataset had already been completed and did not affect any of the methodological choices. The evolutionary conservation was obtained by comparing the independent mice data with the human data from PanglaoDB. The bootstrap frequency of the independent dataset was compared with that of the original dataset using the hypergeometric test.

### Implementation of the algorithm

The pipeline shown in Figure 6 is implemented in our new software DECNEO (DEndrogram of Co-expression with Evolutionary Conservation and Network Enrichment Ordering). We have developed a main and an alternative implementation of DECNEO that are open-source and maintained at https://github.com/sdomanskyi/decneo. Releases are published on the Zenodo archive.

The primary implementation can be installed from PyPI using the command “pip install decneo”. DECNEO modules rely on Python numpy, scipy, pandas, scikit-learn, matplotlib and other packages (see Figure S6 for dependencies). The primary implementation can leverage on a multicore large-memory High-Performance Computing Systems for fast processing of large datasets. The detailed documentation and installation troubleshooting instructions are available at https://decneo.rtfd.io/. The processed data and results reported in this paper are deposited at https://figshare.com/. The source code of the alternative implementation is deposited at https://github.com/sdomanskyi/decneo/validation. The software was optimized for use with the HPCC supercomputer at Michigan State University and also tested on different operating systems and hardware.

### Alternative implementation

The primary implementation of DECNEO was validated by comparing its results to an additional implementation created independently as a software and methodology validation. The alternative implementation has allowed us to verify the robustness of the results under minor differences in the software implementation. Specifically, the differences are: (1) When finding differentially expressed genes for the network enrichment, the alternative implementation filters out downregulated genes and sorts them by p-value. In contrast, the primary implementation sorts genes by the median rank of the t-test statistic. (2) When computing the expression fraction for a set of samples, the median of the expression fraction across samples is taken, instead of the mean across samples as in the main implementation. (3) Lastly, in the alternative implementation, gene measures are not smoothed before merging them into a combination of measures, and the combination of measures is normalized so that the minimum is zero and the maximum is one. (4) In the alternative implementation only endothelial cells, but not non-endothelial cells, are used when generating pseudo samples.

The analysis of endothelial cells in the PanglaoDB-Alona dataset is summarized in Figure S7 and the alternative implementation is shown in Figure S8. These analyses show results similar to Figure 2. Table S6 shows the correlation of bootstrap frequencies between DECNEO main and DECNEO alternative implementation.

### Revealing Multiple Peaks

We observe multiple peaks in the measures and their combinations. These represent the main comberon and other minor comberons. For example Figure S4 shows the “Combination of measures” of *Mus musculus* endothelial cells of PanglaoDB-DCS. We analyzed all the peaks, including the largest one. Each instance of the 100 bootstraps resamples has a set of peaks with the receptors identified in the same way as in Figure S4. We order receptors (vertical axis) and peaks (horizontal axis) according to the “Ward” agglomeration method with Euclidean metric of binary membership, see Figure S9, which is colored according to each peak relative height. The left and top dendrograms highlight receptors and peaks groups. To focus on the most prominent peaks we obtained 15 groups of receptors and 15 groups of peaks. After determining the density of the receptors in each receptor-peak group we show in Table S7 only those with at least 0.5 density. All the groups of peaks that correspond to one group of receptors are aggregated to determine the average frequency and average peak height.

### Tissue-specific receptors evaluation

We have used the analysis pipeline, illustrated in Figure 6, separately on several tissue-specific studies (SRAs) from PanglaoDB annotated by DCS, from *Mus musculus* and *Homo sapiens* datasets. To compare the processed SRAs, we calculated the Pearson correlation of the bootstrap frequencies, shown as a heatmap in Figure 3. The SRAs are ordered according to the Ward [35] agglomerative clustering of the Euclidean similarity metric of the Pearson correlation of bootstrap frequencies.

The top tissue-specific receptors are shown as a heatmap in Figure 4, to show that some receptors can be found in all tissues but others are specific to one or a few tissues. SRA studies of the same tissue were combined by averaging the largest peak bootstrap frequencies. The lists of the top-10 receptors for each of the analyzed tissues and the bootstrap frequency cutoffs are given in Table S8.

### Metascape enrichment analysis

For pathway enrichment analysis, Metascape [44] calculates the p-value defined as the probability from the cumulative hypergeometric distribution and corrects it with the Bonferroni method to give the q-value.

### Ligand-receptor pairs in choroid fibroblasts and endothelial cells

The computational methods used our main software (DECNEO) with an additional set of scripts, also distributed on GitHub as part of the DECNEO project (github.com/sdomanskyi/decneo).

Intercellular communication is analyzed for two cell types at a time. This paper analyzed choroid fibroblast ligands and choroid endothelial receptors, using single-cell RNAseq data from [21,45]. Two-dimensional matrices are built listing ligands on the y axis and receptors in the x axis. Each axis represents one of the dendrograms described in DECNEO and shows the “combination of measures” from that dendrogram. The receptors and ligands are ordered so that co-expressed molecules within single cells are closer to each other, as was done for Figure 2.

For each ligand-receptor pair in the matrix we calculate the number of “known ligand-receptor pairs”, from [26], within their neighbors in the dendrogram. The width of the window used to find neighbors is set to 50. The neighbors are found after a bootstrap procedure with 100 resamples. Only genes that are found in more than 30% of resamples are counted.

The script also calculates the mean of the scaled “combination of measures” from DECNEO for ligands and receptors. The matrix within Figure 5 shows the color-coding of the average of this mean of “combination of measure” for ligand and receptors with the measure of “known ligand-receptor pairs” within the neighbors. Only values above 0.5 are shown in the Figure. The box shown in Figure 5 was obtained using the clustering algorithm DBscan [46].

## ACKNOWLEDGEMENTS

This work was supported by the National Institutes of Health (No. R01GM122085). We are very grateful for the statistical feedback received from Michael G. Walker and Karen Messer. We also wish to thank Christopher Wills for his comments.

## COMPETING INTERESTS

CP and GP own shares of Salgomed, Inc.

## Notes

https://github.com/sdomanskyi/decneo

