## SUPPLEMENT for "Naturally occurring combinations of receptors from single cell transcriptomics in endothelial cells"

### Supplementary Figures

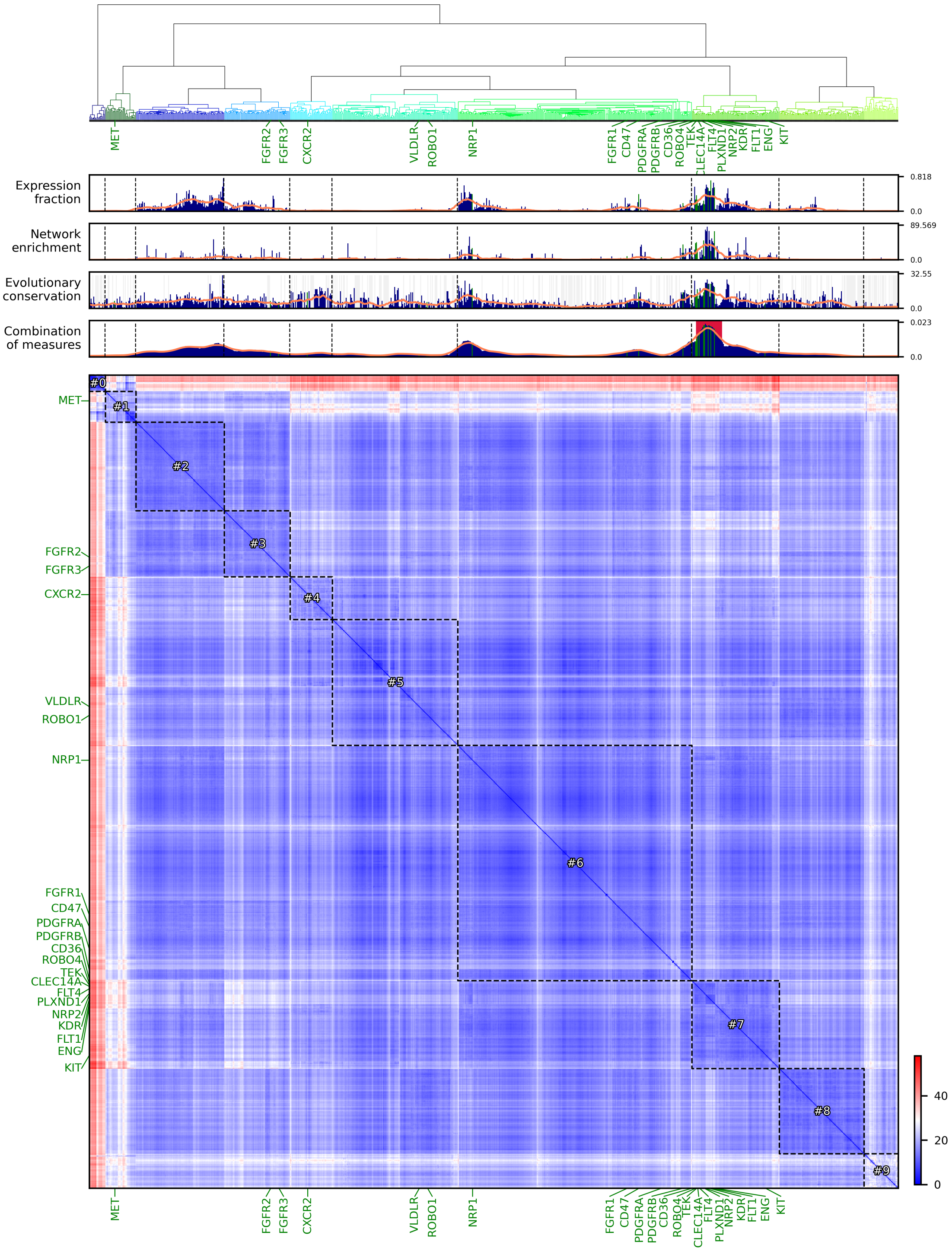


**Figure S1.** Top shows the ordering by single cell co-expression of receptors in endothelial cells of PanglaoDB annotated by DCS for *Mus musculus*. This is identical to Figure 2A. The heatmap shows similarity (Euclidean distance) of correlation of the receptors with all genes arranged according to the optimized dendrogram ordering.

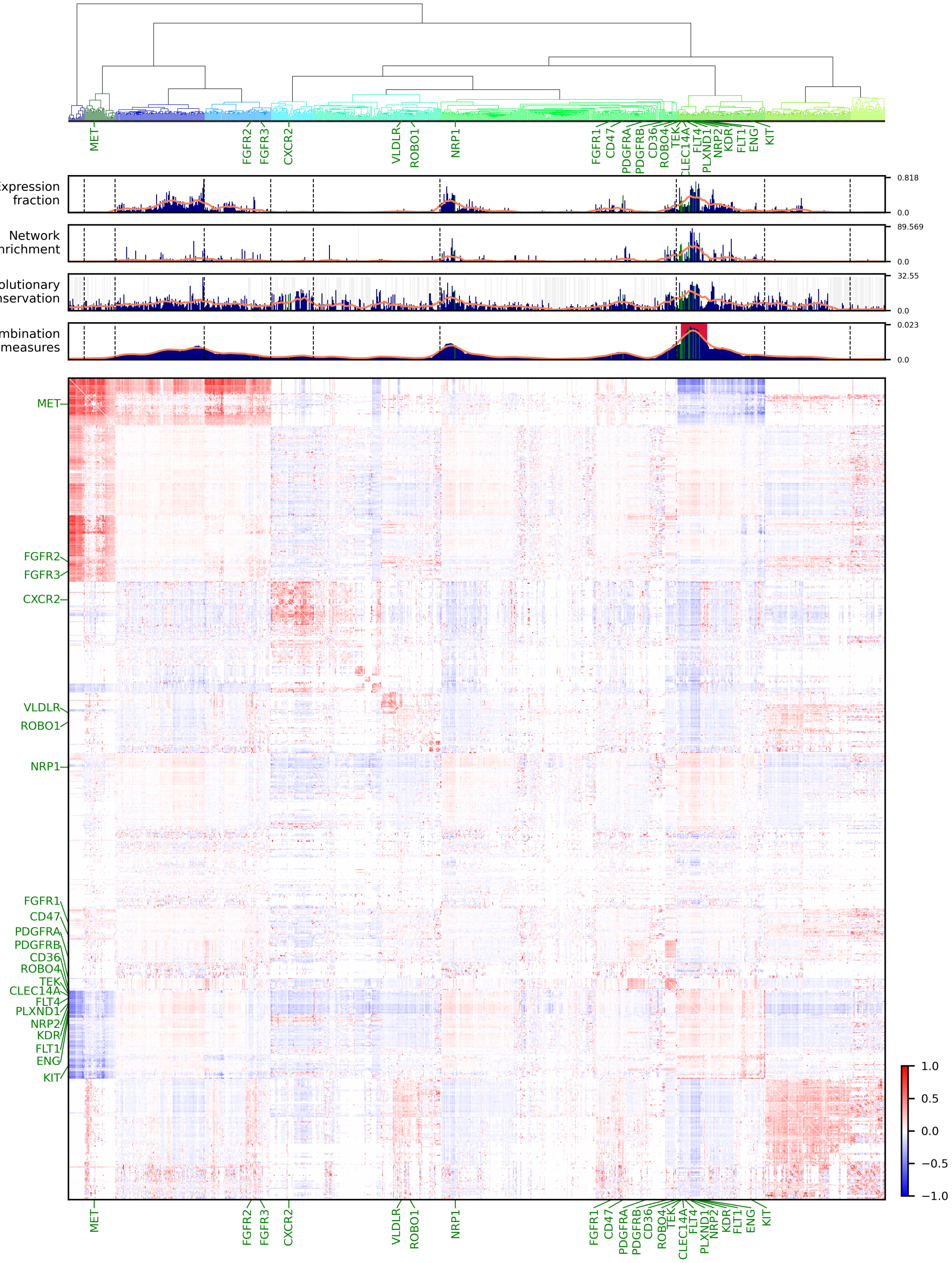


**Figure S2.** Top shows the ordering by single cell co-expression of receptors in endothelial cells of PanglaoDB annotated by DCS for *Mus musculus*. This is identical to Figure 2A. The heatmap shows correlation among the receptors arranged according to the optimized dendrogram ordering. This Figure shows that the correlation among the receptors alone would not be sufficient to detect the comberon (highlighted with a red background).

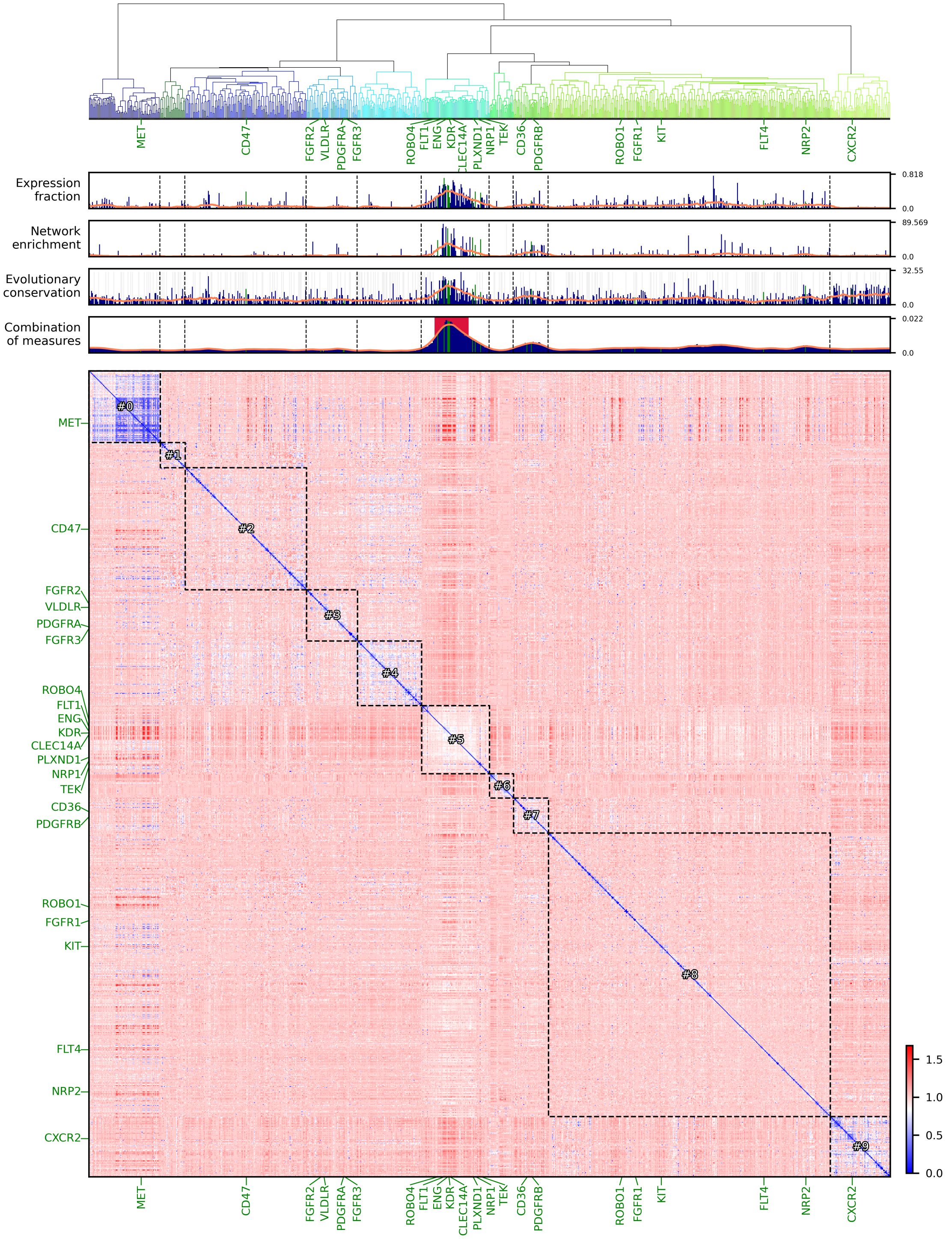


**Figure S3.** Top shows the ordering of the dendrogram by Pearson correlation among receptors in endothelial cells of PanglaoDB annotated by DCS for *Mus musculus*. The heatmap shows Pearson correlation of the receptors arranged according to the optimized dendrogram ordering. The comberon (red background) can be found but the number of known angiogenesis receptors within it is lower compared to the main method (5 vs 10).

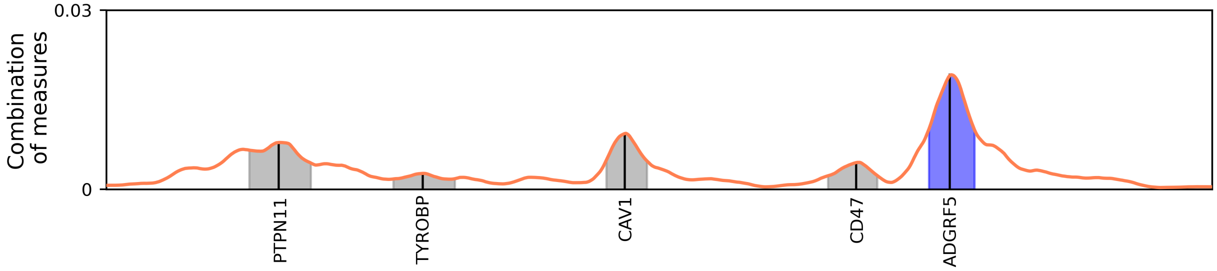


**Figure S4**. Peak gene memberships for endothelial cells from PanglaoDB *Mus musculus* annotated by DCS. “Combination of measures” measures show 5 peaks in this realization. Genes belonging to largest peak are highlighted in blue. The 38 genes included in the largest peak (main comberon) are ADGRF5, ADGRL4, ADRB2, AQP1, CALCRL, CD300LG, CD36, C40, CD74, CD93, CDH5, CLEC14A, CLEC1A, CXCL16, ENG, ESAM, FLT1, FLT4, GPIHBP1, GPR20, IL4R, ITGA4, KDR, LYVE1, NPR1, NRP2, OSMR, PECAM1, PLXND1, PTPN18, PTPRB, RAMP2, ROBO4, SIRPB1, STAB1, TEK, TIE1 and TWF2.

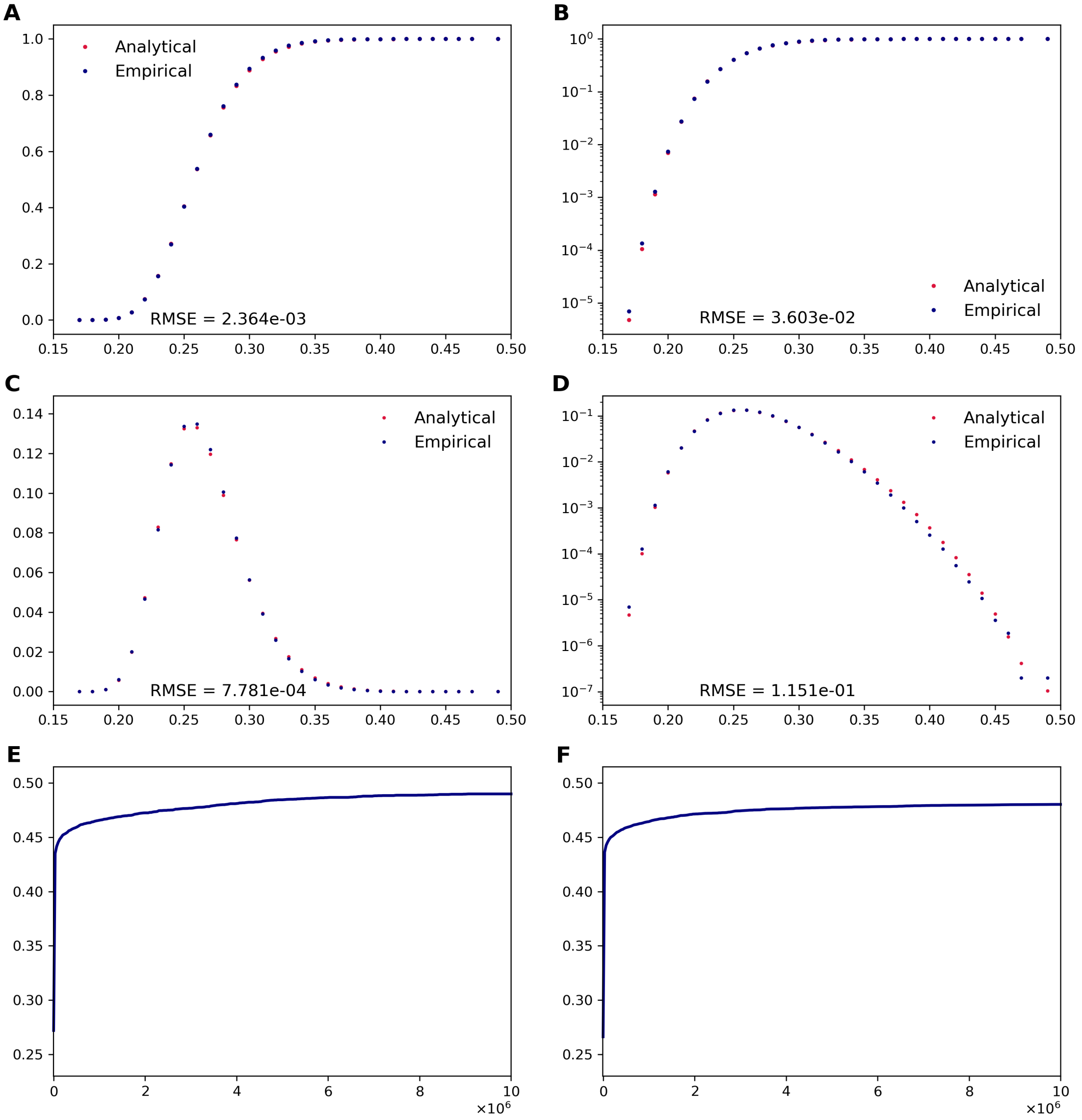


**Figure S5**. Empirical distribution of the maxima of $10^7$ realization of 100 randomizations of the dendrogram order of the "Combination of 3" measure done by the DECNEO main implementation of DCS annotated PanglaoDB Mus musculus datasets. This distribution is binned according to all possible values and presented with blue markers. The generalized extreme value distribution, $F(x;\xi,\mu,\sigma) = \exp\left[-(1-\xi\,(x-\mu)/\sigma)^{1/\xi}\right]$, is found to fit the empirical distribution, where the fitted parameters $\xi = 9.643 \cdot 10^{-2}$, $\mu = 2.523 \cdot 10^{-1}$ and $\sigma = 2.727 \cdot 10^{-2}$ determine the shape, location and scale of the distribution, calculated for the same bins as the empirical distribution and displayed with red markers. The horizontal axis in the first four panel is the maximum frequency in a realization. Panels show (A) cumulative probability distribution function (CDF) in linear scale, (B) CDF in logarithmic scale, (C) discrete probability distribution function (PDF) in linear scale, and (D) PDF in logarithmic scale. The numerical difference between analytical and empirical distribution is obtained by root mean square error (RMSE), indicated in each of the panels. Panel (E) shows the empirical distribution of the maximum values increasing with the number of realizations (horizontal axis). Panel (F) shows the model distribution of the maximum values increasing with the number of realizations (horizontal axis).

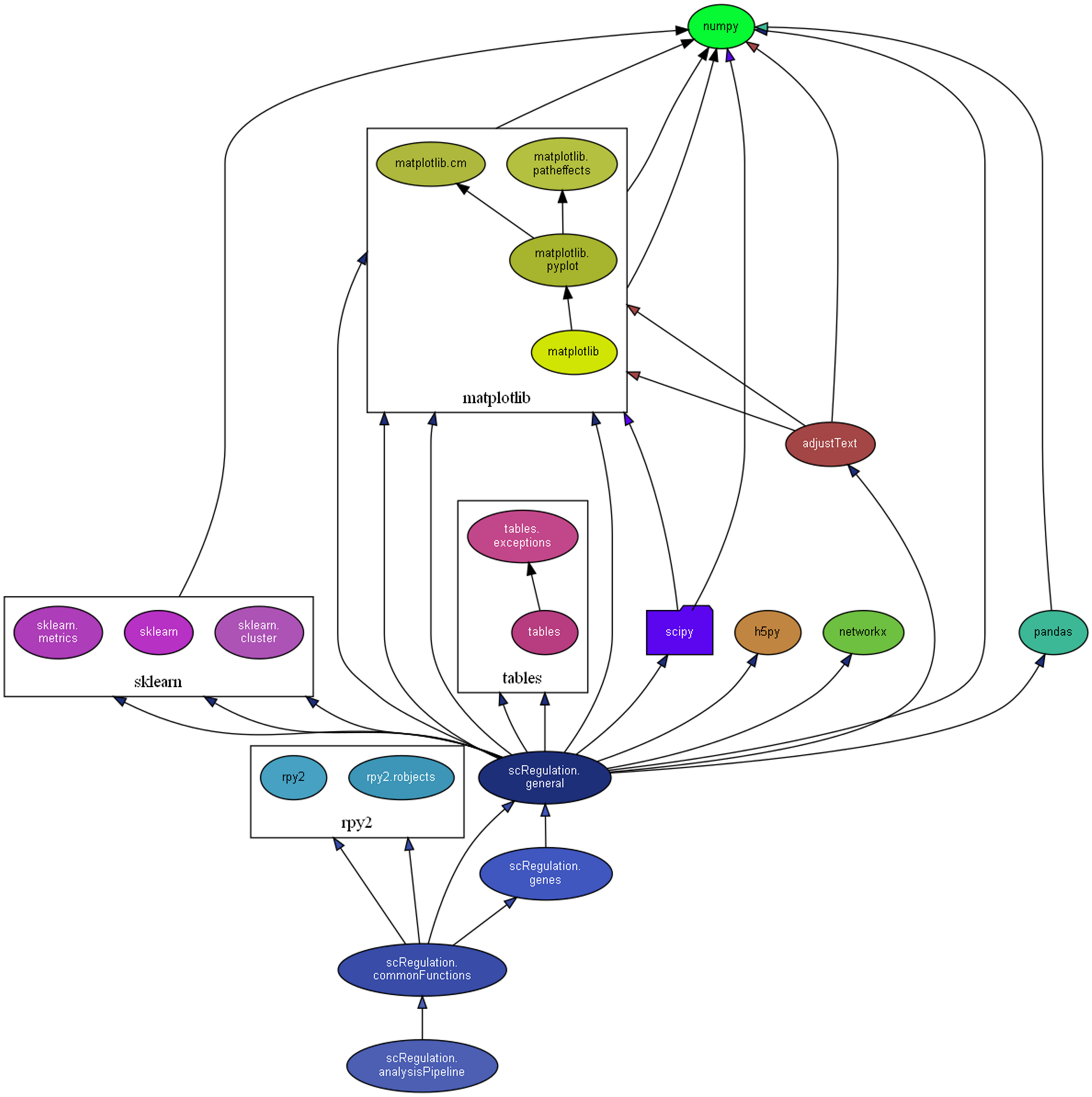


**Figure S6**. Software modules dependency. DECNEO modules are relying on Python-based numpy, scipy, pandas, scikit-learn, matplotlib and other packages.

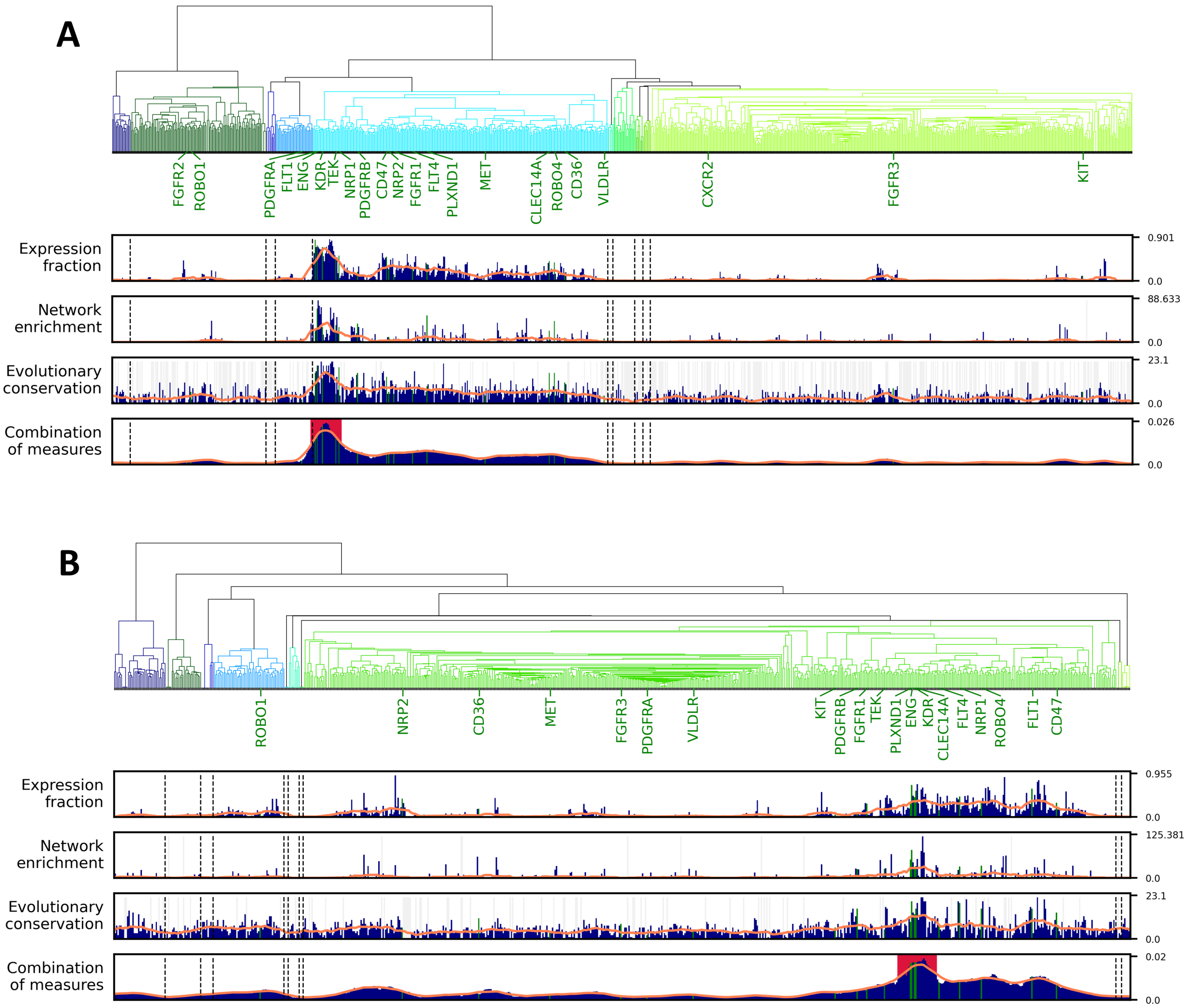


**Figure S7**. Ordering by single cell co-expression of receptors in endothelial cells of PanglaoDB annotated by Alona for: (A) *Mus musculus*, and (B) *Homo sapiens*. See **Figure 2** for detailed description of the dendrogram and information panels.

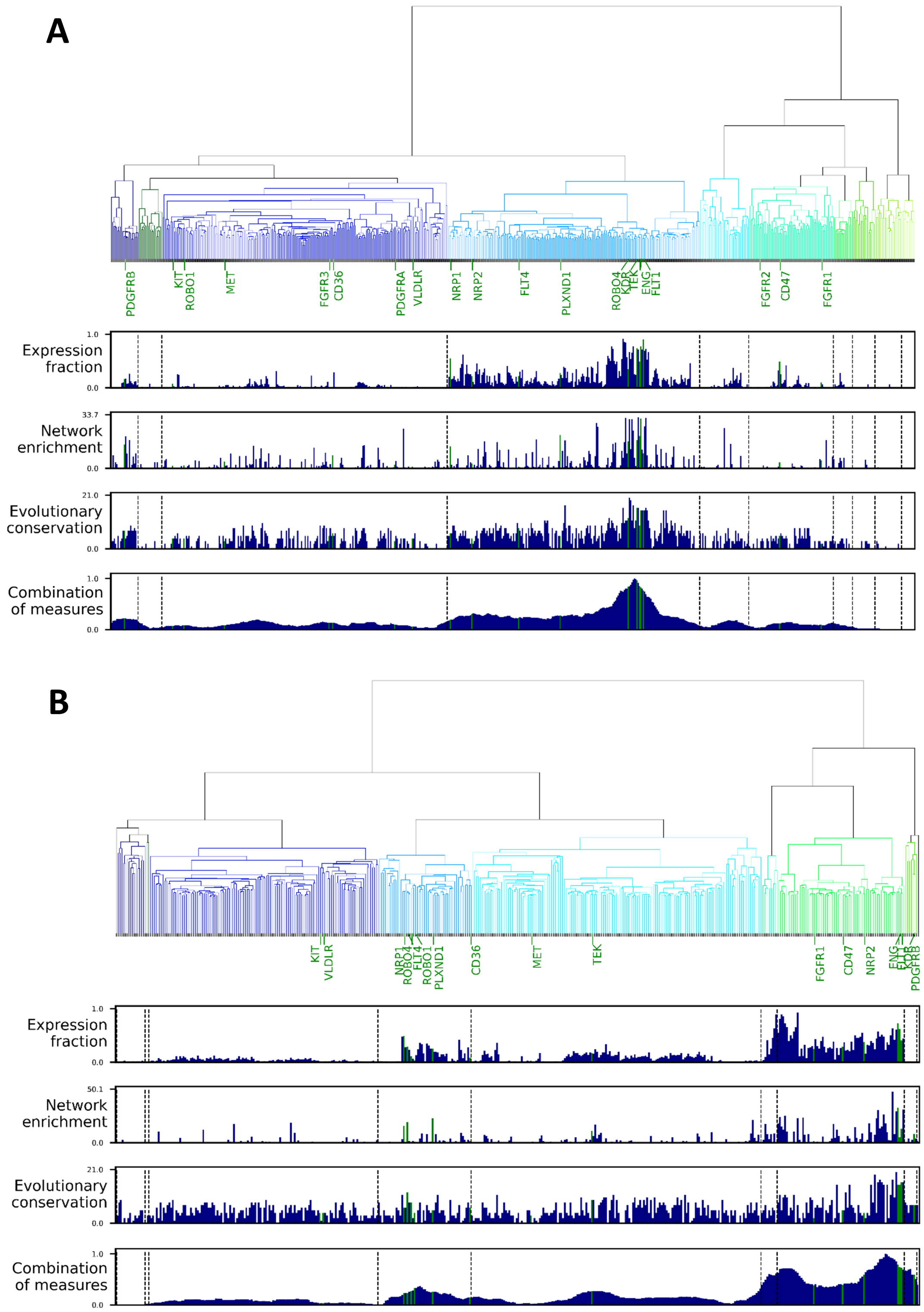


**Figure S8**. Alternative implementation used for validation. PanglaoDB annotated by Alona for: (A) *Mus musculus*, and (B) *Homo sapiens*. See **Figure 2** for more detailed description of the dendrogram and information panels.

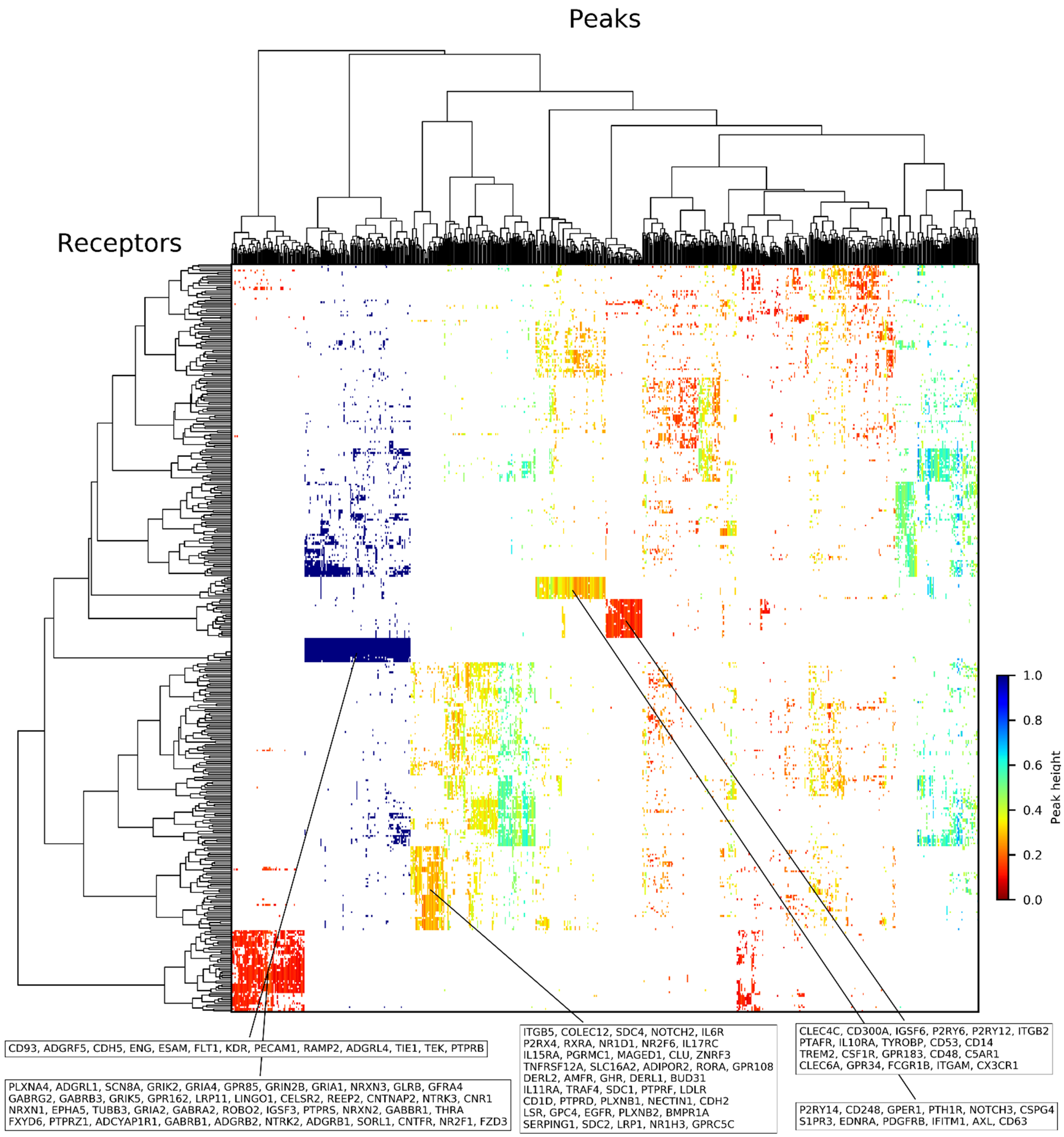


**Figure S9**. Genes from multiple peaks of 100 bootstrap experiments generated from PanglaoDB *Mus musculus* data annotated by DCS and analyzed with DECNEO main implementation for the “Combination of 3” measure. Color-coding indicates peak heights. White color indicates that a gene in not in a given peak. Genes and peaks are ordered by agglomerative clustering with method Ward. Dendrograms highlight genes and peaks groupings. Showcased clusters have their gene memberships annotated in the information boxes.

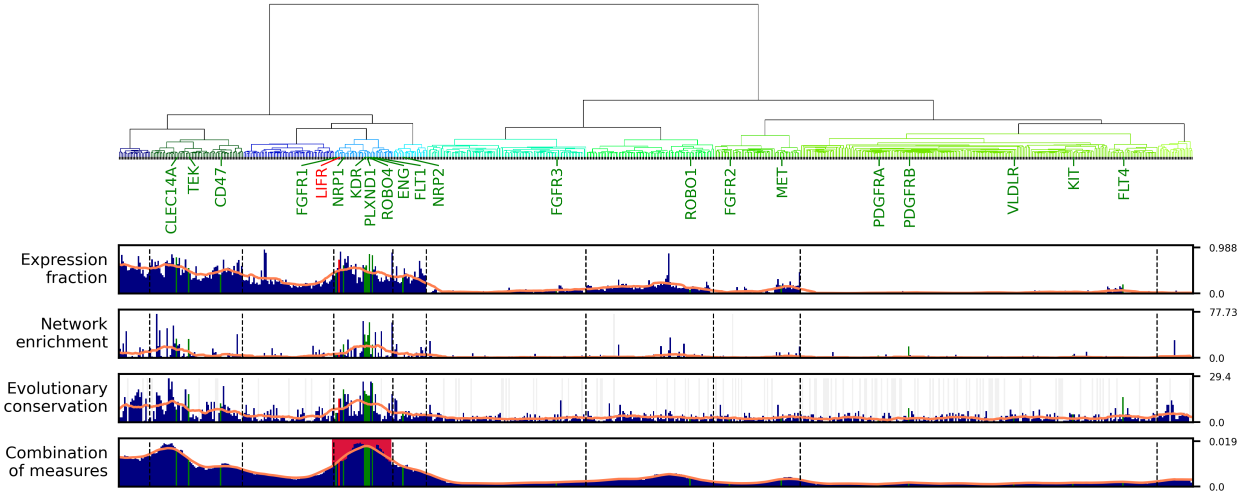


**Figure S10**. Dendrogram ordered by single cell co-expression of receptors in endothelial cells of the choroid dataset. The procedure is similar to that shown in **Figure 2** for the main dataset. The main comberon is shown with a red background. LIFR is highlighted in red.

### Supplementary Tables

**Table S1**. Alternative choices for the gene co-expression metrics, clustering similarity metrics and linkage methods analyzed for PanglaoDB-DCS dataset. See file "Table S1.xlsx".

**Table S2**. Validation of the comberon by splits of the PanglaoDB-DCS *Mus musculus* data. See file "Table S2.xlsx".

**Table S3**. Comberon GO enrichment analysis filtered to show terms from the Biological Process class. See file "Table S3.xlsx".

**Table S4**. Complete version of **Table 1** with bootstrap frequencies for 879 receptors. See file "Table S4.xlsx".

**Table S5**. Independent dataset composition. See file "Table S5.xlsx".

**Table S6**. Pearson correlation of largest-peak bootstrap frequencies from **Table S4**. "Alona (alt.)" is the alternative implementation of DECNEO using the Alona endothelial cell annotation.

| *Approach* | DCS | Alona | Alona (alt.) |
|---|---|---|---|

| | *Species* | *Mus musculus* | *Homo sapiens* | *Mus musculus* | *Homo sapiens* | *Mus musculus* | *Homo sapiens* |
|---|---|---|---|---|---|---|---|
| DCS | *Mus musculus* | 1.00 | 0.76 | 0.85 | 0.55 | 0.64 | 0.58 |
| | *Homo sapiens* | | 1.00 | 0.75 | 0.62 | 0.53 | 0.69 |
| Alona | *Mus musculus* | | | 1.00 | 0.65 | 0.79 | 0.71 |
| | *Homo sapiens* | | | | 1.00 | 0.61 | 0.70 |
| Alona (alt.) | *Mus musculus* | | | | | 1.00 | 0.67 |
| | *Homo sapiens* | | | | | | 1.00 |

**Table S7**. Receptors were clustered from the multiple peak analysis into 11 receptor groups.

| Group | Frequency of group | Average peak height | Number of genes | Genes |
|---|---|---|---|---|
| P1 | 1.00 | 1.00 | 13 | ADGRF5, ADGRL4, CD93, CDH5, ENG, ESAM, FLT1, KDR, PECAM1, PTPRB, RAMP2, TEK, TIE1 |
| P6 | 0.54 | 0.56 | 18 | ADGRL2, APLNR, BAMBI, CAV1, CD151, CD40, CD59, FCGRT, FZD6, IFNGR1, ITGA6, LRP10, LRP8, PROCR, PTPN12, RTP4, STRA6, TNFRSF1A |
| P11 | 0.18 | 0.50 | 29 | ACKR2, ACVRL1, ADORA2A, ADRB2, BMPR2, CCRL2, CLEC2D, DYSF, EPHB4, GPR4, IGF1R, IL10RB, IL2RG, IL6ST, ITGA1, ITGA4, NRP1, OSMR, PEAR1, PLSCR4, PTPN18, PTPRM, SCARF1, SDC3, SIRPB1, STAB1, TGFBR2, TGFBR3, TWF2 |
| P9 | 0.37 | 0.50 | 22 | ADGRE5, ADGRG3, AQP1, CALCRL, CD300LG, CD36, CD55, CLEC14A, CLEC1A, CXCL16, FLT4, GPIHBP1, GPR182, ITGA9, KIT, NPR1, NRP2, PLAUR, PLXND1, RARG, ROBO4, THBD |
| P7 | 0.53 | 0.35 | 13 | ATP6AP2, CANX, CD9, HFE, IFNGR2, LAMP1, LAMP2, LIFR, NCSTN, PPARD, PTPN1, PTPRA, REEP5 |
| P3 | 0.72 | 0.32 | 12 | AXL, CD248, CD63, CSPG4, EDNRA, GPER1, IFITM1, NOTCH3, P2RY14, PDGFRB, PTH1R, S1PR3 |

| | | | | |
|---|---|---|---|---|
| P10 | 0.27 | 0.30 | 45 | ADIPOR2, AMFR, BMPR1A, BUD31, CD1D, CDH2, CLU, COLEC12, DERL1, DERL2, EGFR, GHR, GPC4, GPR108, GPRC5C, IL11RA, IL15RA, IL17RC, IL6R, ITGB5, LDLR, LRP1, LSR, MAGED1, NECTIN1, NOTCH2, NR1D1, NR1H3, NR2F6, P2RX4, PGRMC1, PLXNB1, PLXNB2, PTPRD, PTPRF, RORA, RXRA, SDC1, SDC2, SDC4, SERPING1, SLC16A2, TNFRSF12A, TRAF4, ZNRF3 |
| P2 | 0.79 | 0.27 | 9 | ADIPOR1, APLP2, CD81, CR1, ITGB1, KDELR2, NR3C1, REEP3, S1PR1 |
| P5 | 0.61 | 0.22 | 16 | ABCA1, F11R, FAS, GPR146, IFNAR2, INSR, ITGA5, ITPR2, NR4A1, OCLN, PTPRG, SCARB1, SIGIRR, SLC40A1, TFRC, TNFRSF1B |
| P8 | 0.39 | 0.15 | 21 | C5AR1, CD14, CD300A, CD48, CD53, CLEC4C, CLEC6A, CSF1R, CX3CR1, FCGR1B, GPR183, GPR34, IGSF6, IL10RA, ITGAM, ITGB2, P2RY12, P2RY6, PTAFR, TREM2, TYROBP |
| P4 | 0.68 | 0.13 | 44 | ADCYAP1R1, ADGRB1, ADGRB2, ADGRL1, CELSR2, CNR1, CNTFR, CNTNAP2, EPHA5, FXYD6, FZD3, GABBR1, GABRA2, GABRB1, GABRB3, GABRG2, GFRA4, GLRB, GPR162, GPR85, GRIA1, GRIA2, GRIA4, GRIK2, GRIK5, GRIN2B, IGSF3, LINGO1, LRP11, NR2F1, NRXN1, NRXN2, NRXN3, NTRK2, NTRK3, PLXNA4, PTPRS, PTPRZ1, REEP2, ROBO2, SCN8A, SORL1, THRA, TUBB3 |

**Table S8**. The top-10 tissue-specific receptors of endothelial cells of PanglaoDB and choroid annotated by DCS. The right-most column is the minimum max-peak bootstrap count.

**Table S9**. Analysis of endothelial cells from selected studies (same as in **Figure 4**). Same tissue SRAs are combined into one to find average correlation among them (column “Inside”) and average correlation between them and all other SRAs (column “Outside”). The table is ordered by column “Inside” in descending order. Since most “Inside” values are larger than “Outside” values, we quantitatively observe that the SRAs from similar tissues tend to group together.

| Species | Tissue | Studies | Inside | Outside |
|---|---|---|---|---|
| *Mus musculus* | Hypothalamus | 4 | 0.865 | 0.592 |
| *Mus musculus* | Olfactory bulb | 2 | 0.827 | 0.612 |
| *Mus musculus* | Lung | 5 | 0.817 | 0.579 |
| *Mus musculus* | Subventricular zone | 2 | 0.776 | 0.604 |
| *Homo sapiens* | Testis | 3 | 0.744 | 0.521 |
| *Mus musculus* | Kidney | 3 | 0.734 | 0.594 |
| *Mus musculus* | Bone marrow | 2 | 0.716 | 0.496 |
| *Mus musculus* | Heart | 2 | 0.311 | 0.470 |
